# A complete map of human cytosolic degrons and their relevance for disease

**DOI:** 10.1101/2025.05.10.653233

**Authors:** Vasileios Voutsinos, Kristoffer E. Johansson, Fia B. Larsen, Martin Grønbæk-Thygesen, Nicolas Jonsson, Emma Holm-Olesen, Giulio Tesei, Amelie Stein, Douglas M. Fowler, Kresten Lindorff-Larsen, Rasmus Hartmann-Petersen

## Abstract

Degrons are short protein segments that target proteins for degradation via the ubiquitin-proteasome system and thus ensure timely removal of signaling proteins and clearance of misfolded proteins from the intracellular space. Here, we describe a systematic screen for degrons in the human cytosol. We determine degron potency of >200,000 different 30-residue tiles from more than 5,000 cytosolic human proteins with 99.7% coverage. In total, 19.1% of the tiles function as strong degrons, 30.4% as intermediate degrons, while 50.5% did not display degron properties. The vast majority of the degrons are dependent on the E1 ubiquitin-activating enzyme and the proteasome but independent of autophagy. The results reveal both known and novel degron motifs, both internal as well as at the C-terminus. Mapping the degrons onto protein structures, predicted by AlphaFold2, revealed that most of the degrons are located in buried regions, indicating that they only become active upon unfolding or misfolding. Training of a machine learning model allowed us to probe the degron properties further and predict the cellular abundance of missense variants that operate by forming degrons in exposed and disordered protein regions, thus providing a mechanism of pathogenicity for germline coding variants at such positions.

## Introduction

In eukaryotes, most intracellular proteins are degraded via the ubiquitin-proteasome system (UPS) (1, 2), although with widely different rates. Hence, human proteins display half-lives ranging from a few minutes to several months (3–5). This large range in turnover rates reflects how efficiently the individual proteins are targeted for degradation, and one key discerning feature is the presence and strength of so-called degradation signals or degrons (6). Most degrons that have been characterized are short linear motifs, embedded within the amino acid sequence. In general, degrons mediate direct interaction with E3 ubiquitin-protein ligases that in turn catalyze the conjugation of ubiquitin to the target protein. Once the target protein is ubiquitylated, it is rapidly degraded by the proteasome. Examples of degrons include the KEN-box and phospho-degrons, that regulate timely degradation of various cell cycle regulators and signaling proteins (6–9). The N-end and C-end degrons constitute other well characterized examples (10–13), where specific amino acid residues in the N- and C-terminal regions, respectively, determine the protein turnover rate.

Beyond these more specific degrons, recent studies have explored the properties of the so-called protein quality control (PQC) degrons, which target misfolded proteins for proteasomal degradation. These studies have revealed that—in contrast to the regulatory, N- and C-end degrons—the PQC degrons are typically enriched for hydrophobic residues (14–16) and overlap with chaperone binding regions (17). Accordingly, PQC degrons are expected to be buried within the core of a natively folded protein but become exposed if the structural stability of the protein is reduced, e.g. by environmental stress conditions or by mutation (18). In this manner, structurally destabilized or misfolded proteins are specifically purged from the intracellular space (6, 19). Finally, in parallel to the PQC-linked degradation, ribosomes are also equipped with a quality control system to ensure degradation of proteins derived from aberrant mRNAs. Thus, if ribosomes stall during translation, the nascent polypeptide is elongated by adding a C-terminal alanine (20) (and in yeast also threonine) tail through a noncanonical elongation reaction (21, 22). This so-called C-terminal alanine and threonine (CAT) tailing leads to exposure of lysine residues for ubiquitylation (23), but also operates as a degron in ribosome-associated quality control (20, 24, 25).

Due to the critical role of degrons in regulating protein abundance, characterizing their properties is essential for furthering our understanding of protein evolution, but also have important biotechnological and medical applications. For instance, in protein design and engineering it may be advantageous to eliminate degrons (26, 27), while gene variants linked to hereditary disease or cancer may generate or destroy a degron leading to an altered abundance of the encoded protein (9, 28). For these reasons, there have been multiple attempts at generating sequence-based degron predictors (16, 28, 29).

Here, we describe a systematic degron screen of a 30-residue library composed of more than >200,000 partially overlapping peptides derived from more than 5,000 human cytosolic proteins with near-complete (>99%) coverage. In total, 19.1% of the tiles function as strong degrons and 30.4% as intermediate degrons, that are primarily degraded via the UPS. We investigate degron dependence on sequence features and develop a two-way convolutional neural network model for degron prediction. We apply the model to obtain a mechanistic explanation for loss of function missense variants in exposed and disordered regions and identify pathogenic gene variants that lead to the formation of new degrons.

## Results

### A massively parallel screen for cytosolic degrons

Inspired by previous attempts at characterizing and mapping degrons in both yeast and human cells (7, 11, 14, 15, 17, 18, 30–32), we aimed to map degrons in all proteins in the human cytosol. We selected all protein-coding genes that are localized to the cytosol according to gene ontology (GO) database (33). The open reading frames of these proteins were then divided into 30 residue (90 bp) tiles, each overlapping by 15 residues with the neighboring tiles (**Fig. 1A**). The length of 30 residues was selected on the basis that peptides of this length would be sufficiently long to contain degrons, while too short to harbor stable protein structures. For practical reasons and due to its large size, the library of >200,000 tiles was split into five sublibraries (see Methods). The degron library was fused to the C-terminus of GFP and, after site-specific integration, expressed from a “landing pad” locus in human HEK293T cells (**Fig. 1B**). As the plasmid does not contain a promoter, any plasmids that fail to integrate should not be expressed, while the Bxb1-catalyzed integration in the landing pad leads to single-copy expression of a GFP-fused tile. Since integration of the plasmid at the landing pad blocks the expression of iCasp9, non-recombinant cells can be depleted from the culture by adding AP1903 (Rimiducid) (34, 35) (**Fig. 1B**). To correct for cell-to-cell variations in expression, the integrated plasmid also produces mCherry from an internal ribosomal entry site (IRES) downstream of the GFP fusion. Finally, fluorescence-activated cell sorting (FACS) is used to separate cells into distinct bins based on the GFP:mCherry ratio, followed by DNA sequencing to quantify the frequency of the tiles in each bin (**Fig. 1B**). Sequencing of the generated library revealed that we had successfully managed to measure 212,658 tiles covering 99.7% of the 5,672 cytosolic proteins (**Fig. 1A**).

**Fig. 1.**
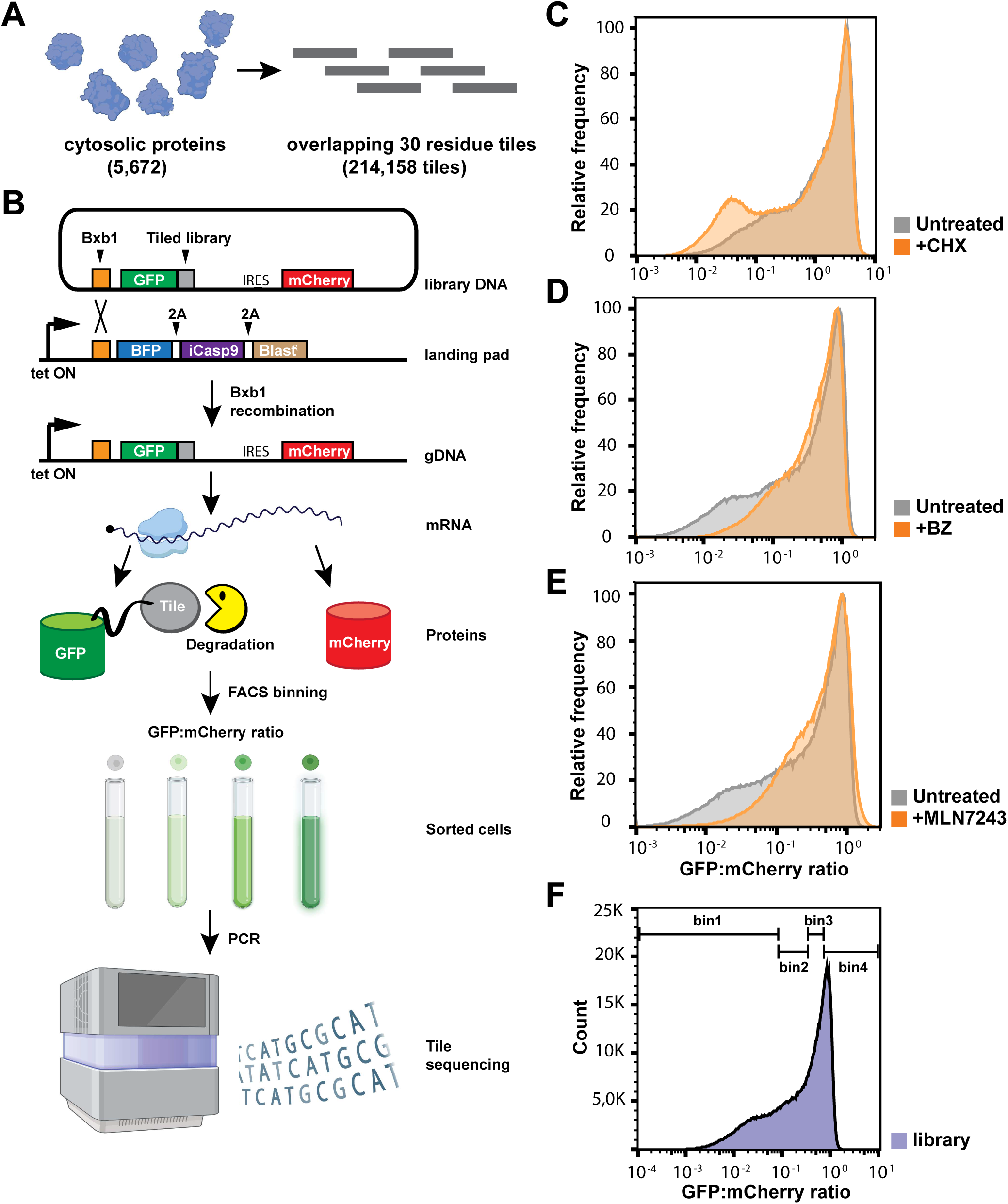
Cytosolic peptide library and degron screening. (A) The peptide library consists of 214,158 different 30-residue partially overlapping fragments covering 5,672 cytosolic human proteins. (B) Schematic illustration of the expression and screening system. The plasmid containing the GFP-fused peptide library also includes an internal ribosomal entry site (IRES), mCherry and a site for Bxb1 recombination into a landing pad in HEK293T cells. Expression from the landing pad is regulated by the Tet-on promoter (bent arrow), that in non-recombined cells drives the expression of BFP, inducible Caspase 9 (iCasp9) and a blasticidin resistance gene (Blast^R^) separated with a parechovirus 2A-like translational stop-start sequence (2A). Upon correct integration, the GFP-fragments and mCherry are expressed from the same mRNA. Using fluorescence-activated cell sorting (FACS) cells are sorted into four equally populated bins and the fragments in each bin can be identified by sequencing. The figure was created with BioRender.com and adapted from (17, 43, 47). Representative distributions of GFP:mCherry ratios in cells expressing the library and either untreated (control) or (C) treated for 8 hours with 10 μg/ml cycloheximide (CHX) (n = 987,000, untreated: n = 932,000), (D) for 16 hours with 15 μM bortezomib (BZ) (n = 887,00, untreated: n = 721,000) (BZ), or (E) 16 hours with 1 μM of MLN7243 (n = 347077, untreated: n = 760,000). (F) A representative flow cytometry profile for cells expressing the library (n=495,000). Bin thresholds used to sort the library into four (1–4) equally populated bins (25% in each bin) are shown as black horizontal bars.

### Degradation is mainly ubiquitin and proteasome dependent

As an initial analysis, the five sublibraries were analyzed by flow cytometry. This revealed that most of the GFP-fused fragments appeared in a large peak representing high GFP:mCherry levels (**Fig. 1C**), indicating that these fragments do not contain strong degrons. However, a large shoulder representing less abundant GFP fusions indicated the presence of degrons. In total the library covered nearly three orders of magnitude in protein abundance as measured by the fluorescence intensity. The flow cytometry profiles of the five sublibraries appeared highly similar (**Fig. S1**).

To test if the low abundance fragments were the result of degradation, the cells were treated with cycloheximide (CHX), which blocks translation. The flow cytometry profile of the treated culture showed a clear shift of the low abundance tiles to even lower GFP:mCherry ratio, indicating that the low steady-state levels of these tiles is a result of degradation (**Fig. 1C**). To further characterize the library, we treated the cells with the proteasome inhibitor bortezomib (BZ), the ubiquitin E1 inhibitor MLN7243, or chloroquine (CQ), which inhibits autophagy. The flow cytometry profiles showed that proteasome and E1 inhibition both clearly shifted the peak of low abundance fragments towards higher GFP:mCherry ratios (**Fig. 1DE**). Conversely, no substantial change was observed with chloroquine (**Fig. S2A**). Treatment with the NEDD8 E1 inhibitor MLN4924 showed a minor stabilization of a limited portion of the low abundance tiles (**Fig. S2B**). Based on these results we conclude that the majority of the low abundance fragments represent degrons targeting the GFP-fusion for ubiquitin-dependent proteasomal degradation, while the contribution from ubiquitin-independent degradation and autophagy is minor, and only a subset of the ubiquitin-proteasome dependent degradation occurs via the NEDD8-dependent cullin-RING ligases, as has been previously reported (32).

### Score distribution and validation

The five sublibraries were separately flow sorted into four bins, each containing 25% of the population (**Fig. 1F**). The landing pad DNA was then amplified by PCR and the tiles were sequenced. Determining the frequency of the tiles in each of the bins allowed us to calculate a degron score for each of the tiles normalized such that a score close to 1 is indicative of high degron potency and a score close to 0 indicates a low degron potency. We performed three biological replicates (separate library transfections) and two FACS replicates for each of the biological replicates. We successfully scored the abundance of 99.7% of the tiles with an average Pearson correlation of 0.98 between pairs of replicate experiments (**Supplementary Data File**).

The degron scores were distributed in four peaks (**Fig. 2A**), and the sequencing showed that peptides in the first peak mainly have read counts in the first FACS bin, and peptides in the second peak mostly in bin 2 and so on. To verify the validity of the obtained scores, we determined the GFP:mCherry ratios for 164 different tiles individually by flow cytometry in low throughput. The results correlated well with the degron scores obtained from the screen (Spearman’s ρ = -0.968) and show a good correlation also within individual peaks (**Fig. 2A**).

**Fig. 2.**
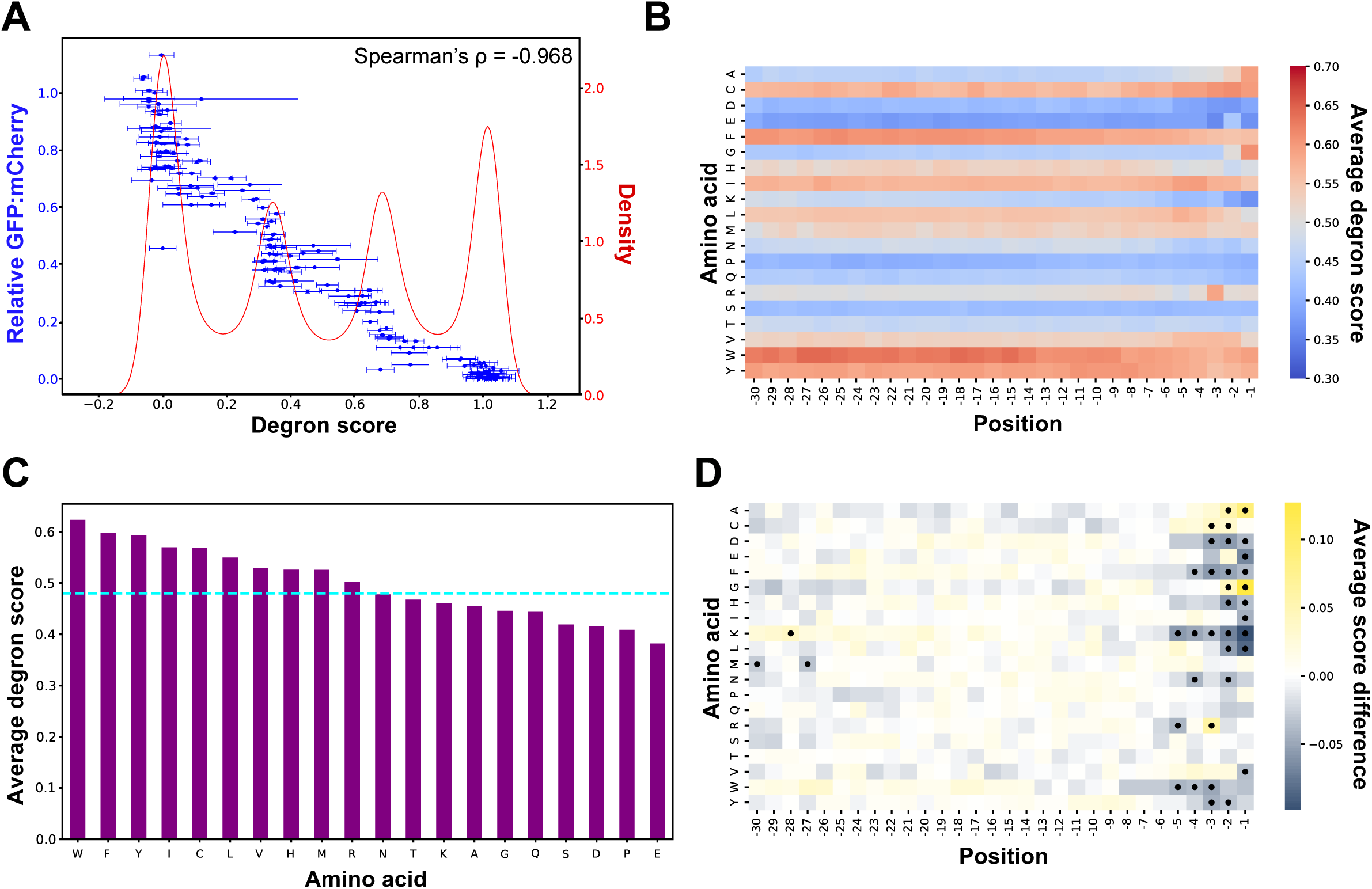
Effects of amino acid residues and position on degron potency. (A) Overlayed degron score distribution (red) with correlation scatter plot comparing the degron scores of 164 random tiles (blue) with their relative GFP:mCherry ratio as measured by flow cytometry and normalized by the GFP:mCherry ratio of the stable tile ENSG00000118898_tile072. Degron score error bars indicate standard deviation and GFP:mCherry error bars indicate standard error. (B) Heatmap showing the averaged degron score of tiles with each amino acid at each position. Red indicates a high degron potency and blue a low degron potency. (C) Bar plot showing the average score of all tiles with each of the amino acids in any of the 25 first positions. The cyan line indicates the total average score of all tiles, which is 0.48. (D) Heatmap showing the difference of the average degron score of exactly one of each amino acid at each position from the average score of exactly one of each amino acid at any other position. Yellow indicates an increase in degron potency at that particular position and blue indicates a decrease. Black dots indicate statistical significance after Bonferroni correction based on Mann-Whitney U Test (p < 0.05/600).

The observation that many peptides mainly have read counts in a single bin is a result of a low level of noise. However, this also confirms that the four-peak distribution is not a property of cellular degradation but a consequence of the sort-seq experiment. Considering the GFP:mCherry fluorescence distribution from flow cytometry a better representative of degradation potency, we numerically transformed the degron scores to this distribution to make an abundance score where zero represents zero fluorescence, *i.e*. low abundance. For the validation point measured in low throughput, this transformation effectively removes the individual peaks in the degron score distribution and recovers the GFP:mCherry fluorescence distribution (**Fig. S3A**). Using the bortezomib treated and untreated library distributions to define degron potency strength, we found in total 19.1% of the tiles function as strong degrons, 30.4% as intermediate degrons, while 50.5% did not display degron properties (**Fig. S3B**). These percentages correspond to abundance score thresholds of approximately 0.04 and 0.22.

### Amino acid composition largely determines degron potency

By calculating the average score of the tiles with a specific amino acid at a certain position, we could determine the general positional effect of that amino acid within a 30-residue tile (**Fig. 2B**). Given that we observed C-end specific effects, we calculated the average degron scores of tiles that contain each of the amino acids within the 25 first positions to determine the general effect that each amino acid has on the abundance of the tiles (**Fig. 2C**). As it has been observed before in human (7, 11) and yeast cells (15), hydrophobic amino acids are generally associated with increased degron potency, with tryptophan, phenylalanine and tyrosine contributing the most to the tile degron potency followed by isoleucine, cysteine, leucine and valine (**Fig. 2C**). The acidic amino acids aspartate and glutamate have a strong stabilizing effect, similar to proline and serine, while glutamine, glycine and alanine are also stabilizing, but to a lesser extent. The general effect of the rest of the amino acids was close to neutral (**Fig. 2BC)**.

Interestingly, in the averaged abundance score map, the effect of previously described C-terminal degrons (7, 11) was also evident. Thus, while both glycine and alanine mainly had a stabilizing effect on the tiles, they considerably increased the degron potency near the C-terminus at positions -1 or -2 (**Fig. 2B**). Similarly, arginine was relatively neutral except at position -3. In general, glutamate was one of the most stabilizing amino acids with a distinct exception at the position -2, most likely due to the C-terminal degron sequences -EE* (**Fig. S4**), -EI*, -EM* and -ES* (asterisk indicating the C-terminus), where the glutamate at the position -2 is critical for the degron recognition (7, 11). All of these cases are well-described C-terminal degrons that are targets of the cullin-RING group of E3 ligases, except for the alanine at positions -1 and -2 and cysteine at positions -1 to -3 (**Fig. 2B**), which have been shown to function as ubiquitin-independent C-terminal degrons (32).

In addition, we found that an alanine dipeptide at the last two positions substantially increased degron potency compared to when it is positioned within the first 10 amino acids of a tile (**Fig. S5**). An increasing number of C-terminal alanine residues showed an increasing degron score, while an increasing number of consecutive alanine residues within the 10 first residues of a tile showed a decreasing degron score (**Fig. S5**). This effect is likely explained by the C-terminal alanine residues resembling the CAT tailing signal associated with ribosome quality control (20–25). Overall, the results showed that in our setup, where tiles were tagged with GFP N-terminally, the degron potency of a tile was largely determined by the amino acid composition, with additional more sequence-specific effects near the C-terminus.

### C-terminal effects are limited to the last five residues

To examine at which positions the C-terminal effects become pronounced, we calculated the average score of all the tiles that have each of the amino acids only once in each position and the equivalent average score of the tiles that have it only once in any other position. We used this approach to control the number of occurrences of the tested amino acid in order to avoid potential biases due to the difference in the representation of the amino acids in our library. This revealed that the C-terminal effects on the degron potency are mostly limited to the last five positions of the tiles (**Fig. 2D**). In addition to the described C-degron effects, this revealed several stabilizing C-terminal sequences. Notably, both aspartate and glutamate at the C-terminus led to a higher protein abundance compared to aspartate and glutamate in the remaining part of the tiles (**Fig. 2D, Fig. S4**). We also observed that with the exception of methionine, cysteine, and valine, all the hydrophobic residues led to reduced degron potency when at the C-terminus of the tiles. A tryptophan dipeptide seemed to be less degron potent when at the C-terminus than when in the first 10 positions (**Fig. S4**). Possibly, this indicates that the negative charge from the C-terminal carboxyl group blocks the otherwise hydrophobic PQC degron signal, in the same way as aspartate and glutamate counteracts degradation in internal degrons. Finally, we noted a strong stabilizing effect of lysine when located at the C-terminal region of the tiles beginning from position -5. Accordingly, we observed a significant decrease in degron potency of lysine dipeptides at the C-terminus versus within the first 10 positions, an observation that also applied to the dipeptide of glutamine (**Fig. S4**). In conclusion, these results confirm the previously observed C-terminal degrons (11, 13) and reveal that certain amino acids may lead to increased protein abundance when located within the last five positions of the tiles.

### C-terminal lysine residues counter degrons and stabilize short-lived proteins

To further investigate the stabilizing effect of lysine residues near the C-terminus, we selected a range of degrons and analyzed the impact of adding two C-terminal lysine residues. Tile 22 of PABPC1L is an -EE* C-terminal degron, while tile 18 of RIC1 is a -GA* C-terminal degron. Given that histidine is the one of amino acids that overall has the smallest effect when at the C-terminus (**Fig. 2B**, **Fig. S6**), we first added histidine to the C-terminus of these sequences to disrupt the C-degrons. As expected, the abundance levels increased for both C-degrons, while the addition of a lysine dipeptide at the C-terminus of the tile resulted in an even more prominent increase in abundance (**Fig. S7AB**). Next, we tested whether C-terminal lysines can stabilize internal degrons. We added a lysine dipeptide to the C-terminus of the well described so-called APPY degron (17) (**Fig. S7C**). It has been found that the main motif responsible for the degradation of APPY is the RLLL sequence located centrally in the peptide and mutation of the RLLL into DAAA significantly reduces its degron potency (17). Addition of the C-terminal lysine dipeptide markedly stabilized the APPY degron, by approximately as much as mutation to DAAA. Moreover, a C-terminal dilysine further increased the abundance even of the stable DAAA variant (**Fig. S7C**).

Following the logic that amino acids that lower the degron score of a tile can have a stabilizing effect, we wanted to test whether non-degron tiles in our library can increase the abundance of unstable full-length proteins. For that purpose, we used the unstable C152W variant of the protein ASPA (36), to which we C-terminally attached one of the lowest scoring tiles in our library (score = 0.12): tile 125 of CENPF, which is highly acidic and with a lysine at position -3. As expected, ASPA C152W had a clearly reduced abundance compared to wild-type ASPA. Fusion of the CENPF tile to the ASPA C-terminus increased ASPA C152W protein abundance (**Fig. S7D**), and the effect was even more pronounced when two CENPF tiles were fused in tandem (**Fig. S7D**). We conclude that fusion to low degron potency peptides can result in increasing the abundance of various targets, and these stabilizing peptides thus appear to be transferrable.

### Degron scores of previously characterized degrons

To test if any previously characterized degron motifs were captured in our screen, we compared our results with a recently compiled dataset of 678 characterized human degrons (37). The highest average degron scores from our library corresponded to the internal degrons (**Fig. S8**). We also observed that many of the described N-end degrons had a degron score above the mean score of 0.48 in our library. Since our library is positioned at the C-terminus, capturing some of the N-end degrons is likely due to hydrophobicity of N-end degrons. Intriguingly, the previously characterized internal degron motifs which were also included in our library showed a large range in average degron scores. In general, previously characterized degrons that were captured in our screen could all be explained based on their composition rather than their specific sequence motifs. Thus, previously characterized degrons such as the p53 family degron recognized by the MDM2 SWIB domain (38) (F[^P]{3}W[^P]{2,3}[VIL]) that displayed high degron scores in our screen displayed a large average hydrophobicity. However, several previously reported degrons, such as the motif D(S)G.{2,3}([ST]), recognized by SCF-TRCP1, and the motif [AVP].{1}[ST][ST][ST], recognized by SPOP (39, 40), were scored as very poor degrons in our dataset. Possibly these degrons are only active in specific cell types or are dependent on post-translational modifications. However, they may also be dependent on being embedded within certain sequence and/or structural contexts that are not accounted for in our library. In that case, the motifs are not transferrable and thus not strictly degrons.

### Most degrons are located in buried and structured protein regions

To determine how the low and high scoring tiles are positioned in the structure of full-length proteins, we used the average relative accessible surface area (rASA), and the average predicted local distance difference test (pLDDT) from AlphaFold as measures of exposure and how structured the tile is within its native context, respectively. We found that most of the tiles with high degron potency tended to be in structured (high pLDDT) and buried (low rASA) regions of the proteins, in contrast to the more stable tiles of our library that in general were found in exposed (high rASA) and intrinsically disordered (low pLDDT) regions (**Fig. 3AB**). This is in accordance with previous findings (17) and suggests that most of the identified degrons are PQC degrons that only become relevant when they are exposed e.g. upon protein unfolding or misfolding (17, 18).

**Fig. 3.**
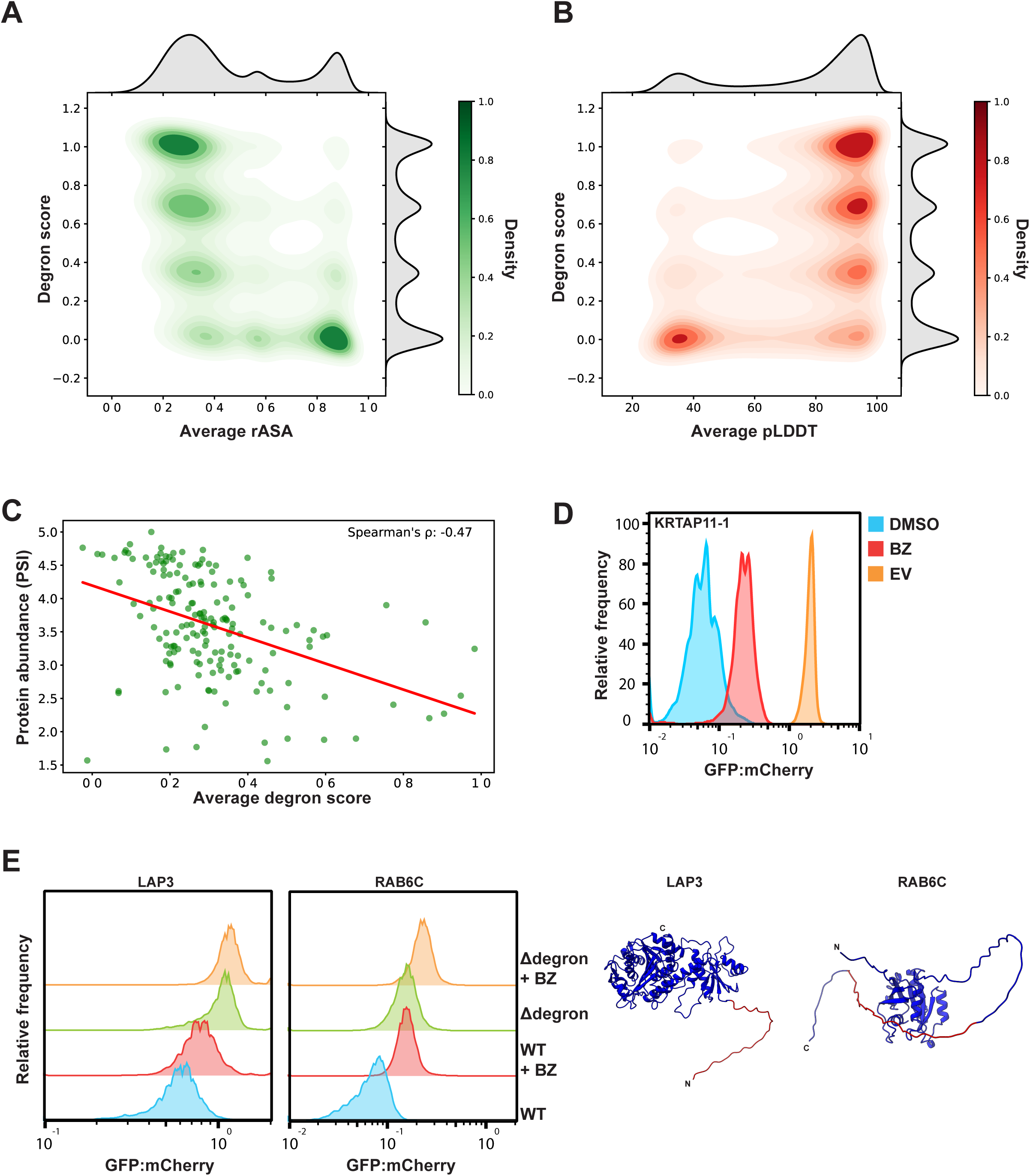
Importance of structural context and exposure of degrons. (A) Kernel density estimate (KDE)plot correlating the average rASA of each tile with its degron score. Dark green indicates high density. (B) KDE plot of the average pLDDT of each tile with its degron score. Dark red indicates high density. KDE was computed with bandwidth = 0.1316. (C) Correlation of average degron score of all tiles of a protein with its protein stability index (PSI) as determined in (11). The correlation is shown only for 173 proteins with the highest average exposure (rASA > 0.7) (D) Representative flow cytometry profile of full-length KRTAP11-1 containing several exposed degrons with BZ (15 μM for 16 h) or without (DMSO) treatment. (KRTAP11-1: n = 1,887, KRTAP11-1 + BZ: n = 1,234). The GFP:mCherry profile of the empty vector (EV) control is shown for comparison (n = 1,730). (E) Representative flow cytometry profiles of LAP3 and RAB6C with BZ (15 μM for 16 h) or without (DMSO) treatment and with degron deleted (Δdegron) or wild type (WT). (LAP3 WT: n = 10,087, LAP3 WT + BZ: n = 7,122, LAP3 DD: n = 7,177, LAP3 DD + BZ: n = 6,548, RAB6C WT: n = 7,499, RAB6C WT + BZ: n = 5,942, RAB6C DD: n = 7,700, RAB6C DD + BZ: n = 5,955). The AlphaFold predicted structures of LAP3 (AF-P28838-F1) and RAB6C (AF-Q9H0N0-F1) are shown on the right with the deleted degrons marked in red. The degron scores of the deleted degrons (shown in red) were as follows: LAP3 tile 1: degron score = 1, average tile rASA = 0.93. RAB6C tile 15: degron score = 0.7, average tile rASA = 0.86. All flow cytometry experiments were performed in duplicate.

### Average degron score predicts the abundance of proteins with highly exposed degrons

In order to investigate the effect of the overall degron potency of a protein on its abundance we attempted to correlate the average degron score of all tiles in a protein with the previously determined abundance of the full-length proteins (11). The protein abundance was expressed as protein stability index (PSI), ranging from 1 to 5 with 1 being low and 5 being high abundance. Overall, the correlation was poor (**Fig. S9**) when considering all the 2,375 different proteins for which a PSI score was assigned. However, when specifically searching for exposed degrons (proteins with an average tile rASA > 0.7), the average degron scores correlated with reduced protein abundance (**Fig. 3C**), and this correlation improved by increasing the rASA threshold (**Fig. S9**). Accordingly, when we experimentally tested the highly disordered keratin associated protein KRTAP11-1 with several exposed degrons (**Fig. S10**), we found that it was rapidly degraded by the proteasome (**Fig. 3D**). To further test the effect of exposed degrons, we selected two proteins, LAP3 and RAB6C, each containing exposed tiles with high degron scores. In both cases, the degradation of the proteins was proteasome dependent and deletion of an exposed degron tile (Δdegron) led to increased cellular levels (**Fig. 3E**). In conclusion, proteins with exposed degrons tend to be short-lived proteasome targets.

### Creating a degron prediction model

Next, we aimed to create a sequence-based model to predict degrons. We used the transformed abundance scores in our modeling to predict GFP:mCherry fluorescence, rather than directly predicting the result of the sort-seq experiment. We first performed a linear regression analysis to identify sequence motifs with predictive power. Based on this analysis, we developed a convolutional neural network with the ability to evaluate peptides of different lengths and with a separate output for C-degron effects.

The linear regression analysis uses lasso regularization to remove features probing less important motifs and thus estimate both the importance and degron potency of sequence motifs. In this analysis we considered simple sequence features including all single amino acids and pairs of amino acids at specific positions and triplets in the C-terminal positions (see methods). The analysis confirmed that the overall amino acid composition is the most important feature in predicting the score with the negatively charged (Glu, Asp) and hydrophobic amino acids (Leu, Phe, Ile, Tyr, Trp, Val, Cys, Met) being most important (**Fig. S11**). The composition effect (position independent) of Lys, Ala and Thr was found to be unimportant, although the three amino acids do have several position-specific effects. As discussed above, this is most notable for Lys that shows a stabilizing effect in the C-terminal positions and gradually less when positioned further from the C-terminus. Position-specific motifs are localized in the C-terminus and interpreted as C-degrons. Our analysis reveals that the strongest C-degrons are the well-known pairs, -EE*, -GG*, -AA*, -GA*, -RxxG*, -RG* and -PG*, but substantial effects are also found for less studied pairs like -KG*, -RxP*, -KxR*, -NEx* and the triplet -PxPA*. C-degron effects are in general stronger towards the C-terminus, notably the gradually changing effect of Lys and -RxxG* which is stronger than -RxxGx* which is again stronger than - RxxGxx*, in agreement with the structure of the CRL2^APPBP2^ ubiquitin-ligase that targets these degrons (41).

Based on the linear regression analysis we next developed a two-channel convolutional neural network (CNN) that we named Peptide Abundance Predictor (PAP) (see Methods) (**Fig. 4A**). One channel consists of a convolutional filter that slides along the peptide to model the composition effect, followed by a global pooling layer that allows evaluation of peptides of different lengths. The second channel considers position-specific effects in the C-terminal positions. Compared to the linear regression model, the CNN can capture non-linear effects and motifs involving any number of amino acids if these are within the considered window of positions (**Fig. 4B**). Thus, PAP efficiently captures both internal and C-terminal degrons (**Fig. 4B**). We tested PAP on the 164 low-throughput validation tile measurements and found that it successfully predicted their abundance (Pearson’s r = 0.85) (**Fig. S12**). An online accessible version of PAP is available via GitHub and Colab.

**Fig. 4.**
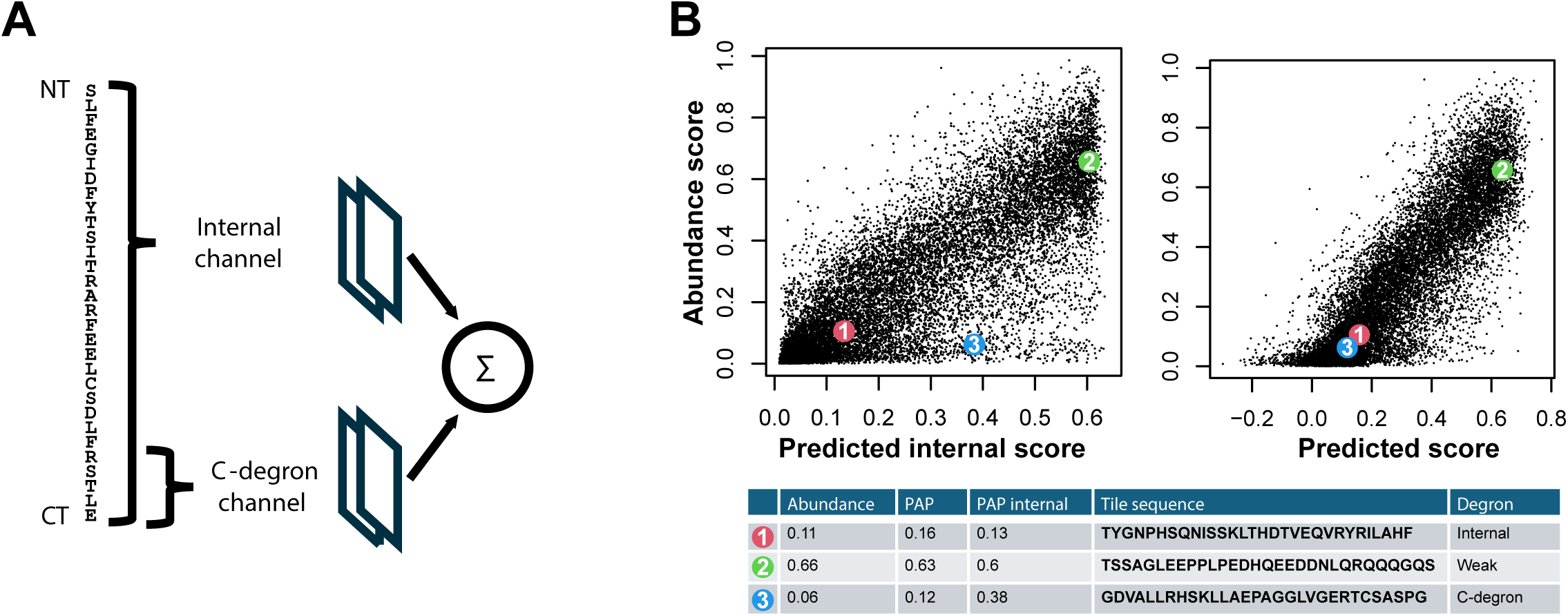
Peptide abundance predictor. (A) Architecture of the two-way convolutional neural network. The internal channel is composed by a convolutional filter followed by global pooling and a few dense layers. The C-degron channel uses a one-hot encoding of the last C-terminal positions also followed by a few dense layers. (B) Predicted scores of the internal channel only (left) and the full model (internal plus C-degron; right) versus the measured abundance scores of the holdout test tiles (Pearson 0.82 and 0.87 respectively). Three tiles are highlighted as examples of a composition driven degron (1), a high abundance tile (2) and a tile with a C-degron (3). The sequence and scores of the highlighted tiles are shown in the table.

### Degron predictions on exposed and unstructured regions correlate with protein abundance

To assess the effectiveness of the predictor, we tested to what extent it could capture the abundance of full-length protein variants in human cells. Specifically, we first collected protein abundance data from deep mutational scanning on eight different proteins ASPA (42), CYP2C19 (43), CYP2C9 (44), NUDT15 (45), PRKN (46), VKOR (47), PTEN and TPMT (34). The difference in PAP-score between a single amino acid substitution variant and the WT (ΔPAP) was calculated with the C-degron term only applied to the C-terminal tile to describe the full-length context. Then, ΔPAP was correlated with the equivalent protein abundance scores within a sliding window of five positions for all the positions in each protein. This revealed a significant difference in the Pearson correlation coefficient distributions between exposed (rASA > 0.7) and less exposed (rASA ≤ 0.7) residues (**Fig. S13**), again suggesting that the direct relevance of degrons in full-length proteins is confined to exposed regions. Thus, the regions showing strong positive correlation are highly exposed and often unstructured, while structured and buried regions show low or negative correlation (**Fig. 5**). The negative correlations are likely a consequence of the predominantly hydrophobic composition of buried and structured regions.

**Fig. 5.**
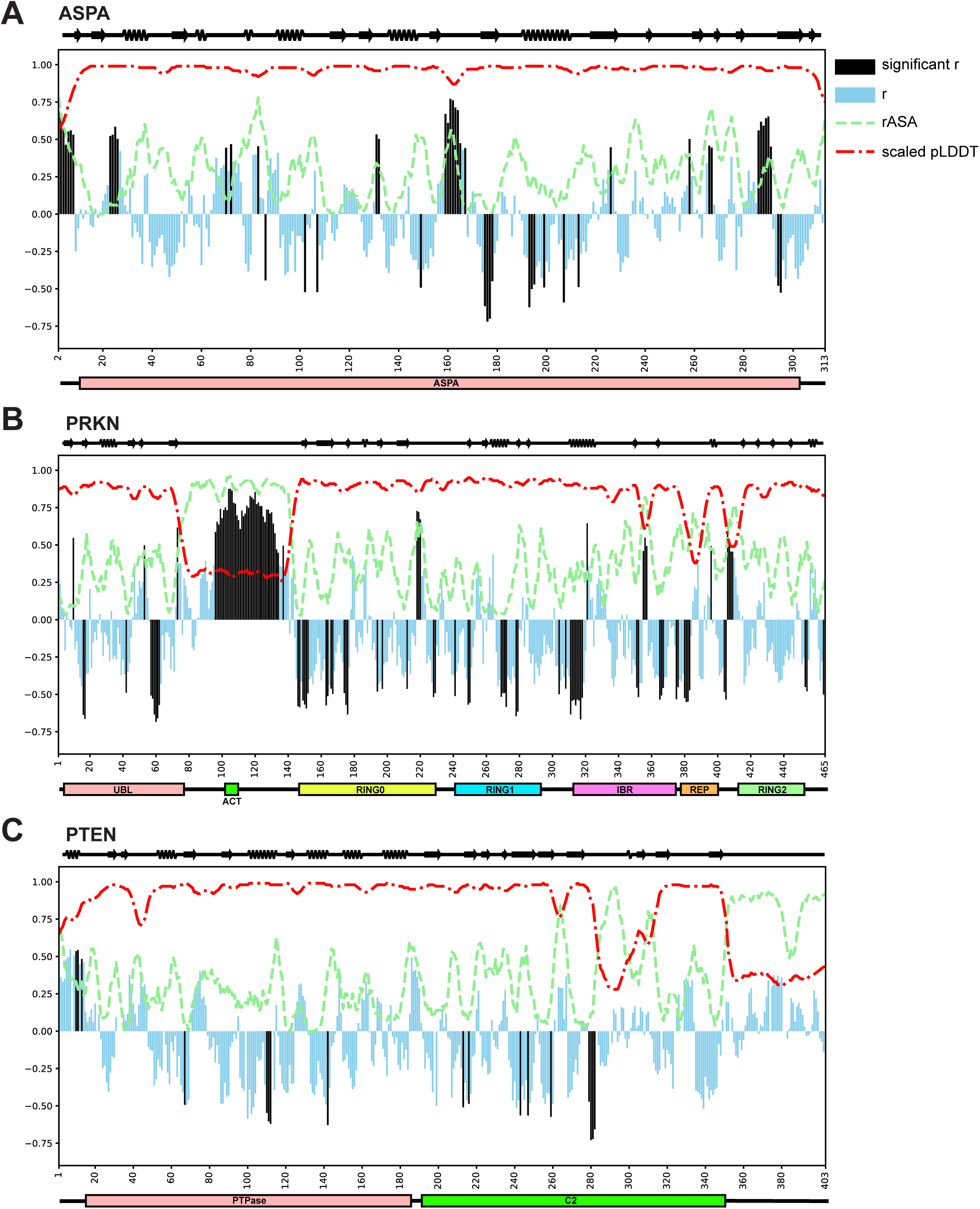
ΔPAP can predict the abundance of missense protein variants. Correlation map ΔPAP against abundance score of single amino acid substitution variants of (A) ASPA, (B) PRKN (Parkin) and (C) PTEN. Bars show the Pearson correlation coefficient of all the scored variants against their predicted ΔPAP for a sliding window of five residues, with the coefficient value assigned to the central residue of the five. The significance of the correlation for every five-residue window was assessed by calculating a p-value. Black bars indicate statistically significant Pearson correlation coefficients with p < 0.05/m, where m is the number of tests conducted for each protein. The average rASA (green line) and pLDDT (red line) of each residue window are also shown. The x axes indicate the amino acid position in each protein. The secondary structure and domain composition of each protein are shown above and below each plot, respectively.

Introducing hydrophilic residues in buried degrons reduces their degron potency and therefore increases the PAP score but results in unfolding and subsequent degradation of the protein, potentially through exposing neighboring intrinsic degrons. However, for the exposed regions, degron potency predictions could help explain both decrease and increase in full-length protein abundance. We experimentally tested this by introducing two mutations that decrease the degron potency in an exposed region of LAP3. In both cases we observed a robust stabilization of the protein (**Fig. S14**). For ASPA, the unstructured N-terminal region (windows centered at positions 2–8) showed a strong correlation between ΔPAP and protein abundance scores, similar to other exposed regions (**Fig. 5A**). For the region covering residues 159–166, we experimentally tested the correlation by fusing that eight-residue long tile to the C-terminus of GFP, followed by five consecutive histidine residues to avoid any potential C-degron effects. We then introduced single-amino acid substitutions and measured the GFP:mCherry ratio for each of them. The ratios correlated (Pearson’s r = 0.85) with the abundance scores of the equivalent protein variants (**Fig. S15**), revealing that at this position ΔPAP successfully predicts full-length protein abundance. An interesting area with a correlation is in the five-residue window centered at position 258. There, we observed a positive correlation for all the single amino acid substitution variants except for substitutions from the prolines at positions 257 and 260, which if mutated profoundly destabilize the protein (**Fig. S16**), perhaps indicating a critical loop structure that is disrupted by substitution of the prolines.

As expected in PRKN, the highly exposed disordered region that lies between UBL and RING0 (residues 96–132) is characterized by a positive correlation between ΔPAP and abundance score (**Fig 5B**). We have previously experimentally demonstrated the correlation between degron potency and abundance in the center of this disordered region (46). Other regions of note include the residues flanking the positions 218, 356 and 407 which all correspond to mostly unstructured exposed loops. In PTEN we found a ΔPAP-abundance score correlation within the exposed N-terminal region (**Fig 5C**). Surprisingly, the highly exposed and disordered C-terminus of the protein did not show a particularly strong correlation. These results could indicate that even though according to AlphaFold the C-terminus is disordered and solvent exposed it might be inaccessible when the protein is in its inactive state.

Data for the other proteins are included in the supplementary material (**Fig. S17**) and reveal similar trends. In conclusion, ΔPAP scores explain some, but not all, of the full-length protein abundance effects for variants at exposed positions.

### PAP can provide mechanistic insight into disease associated missense variants

Given the success of ΔPAP in explaining full-length protein abundance for variants in disordered and exposed regions, we proceeded to investigate if any disease-linked missense protein variants are pathogenic because they cause the formation of novel degrons. Comparison of the ΔPAP for pathogenic and likely pathogenic missense variants with the ΔPAP of benign and likely benign variants within exposed regions (average rASA > 0.7) revealed a significantly (p = 5.4 x 10^-6^) lower ΔPAP for the pathogenic/likely pathogenic mutations, indicating that *de novo* degron formation might be a cause of pathogenicity of some of them (**Fig. 6A**).

**Fig. 6.**
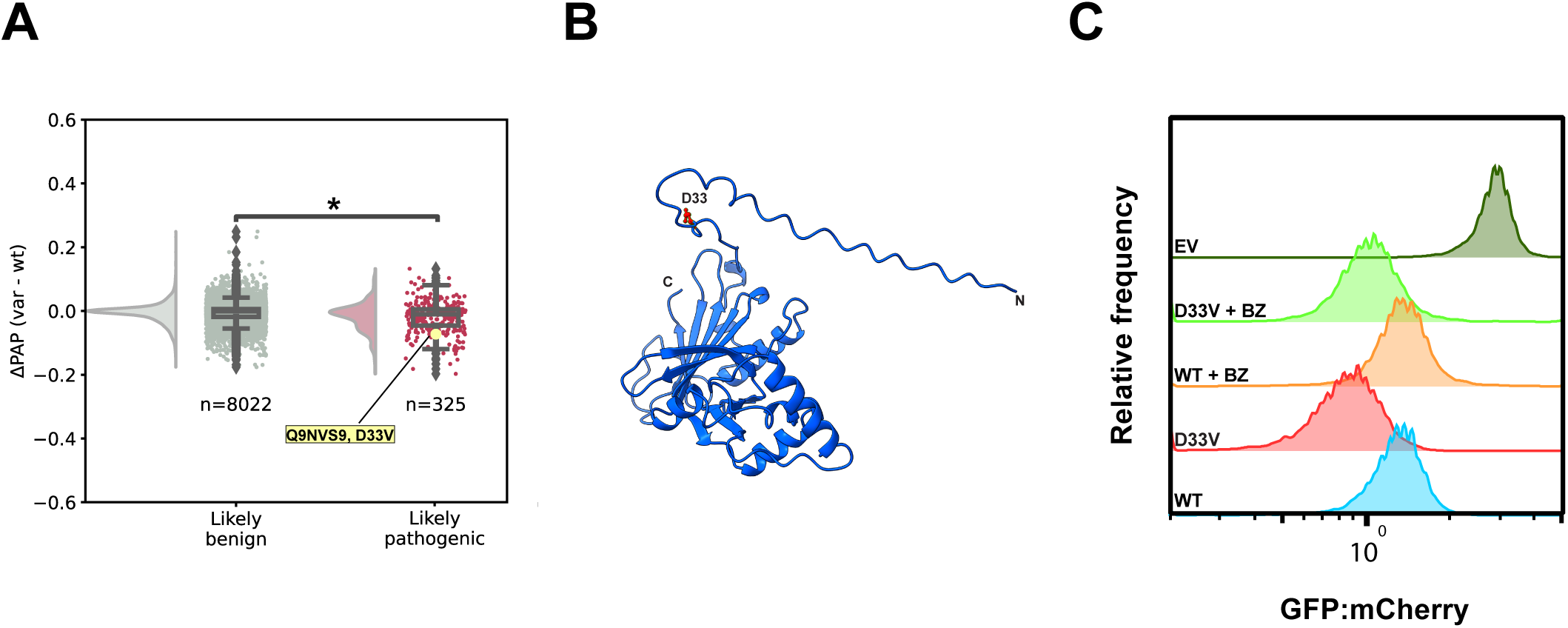
PAP can detect potential pathogenic *de novo* degron creation from missense mutations. (A) ΔPAP of all missense variants within exposed regions (average rASA ≥ 0.7, window size = 5). The central line is at the median of the two populations and the boxes indicate the interquartile range (IQR). The whiskers show the data range within 1.5x the IQR, diamond shaped data point are outliers. The number of data points (n) is shown in the plot. The asterisk indicates statistical significance in a Kruskal-Wallis test (p = 5.4 x 10^-6^) (B) PNPO AlphaFold predicted structure (AF-Q9NVS9-F1). The D33 is shown in red and the N- and C-terminus of the protein are annotated. (C) Representative FACS profiles of PNPO WT and D33V with BZ and without (DMSO) treatment (15 μM for 16 h) (WT: n = 9,933, D33V: n = 10,096, WT + BZ: n = 7,588, D33V + BZ: n = 7,619, EV). An empty vector (EV) control was included for comparisons.

To further test this experimentally, we selected the D33V variant of the pyridoxine-5’-phosphate oxidase (PNPO) enzyme, which has been linked to the rare autosomal recessive disease known as PNPO deficiency (MIM: 610090), which manifests as neonatal epileptic encephalopathy. The N-terminal region, including D33, is highly solvent exposed (**Fig. 6B**) and *in vitro* studies have shown the D33V variant to be enzymatically active and structurally stable (48). In agreement with the ΔPAP of -0.073, we observed a clear proteasome-dependent decrease of PNPO D33V abundance (**Fig. 6C**), suggesting that the D33V variant operates by creating an exposed quality control degron, that in turn results in insufficient cellular levels of the otherwise functional enzyme.

In conclusion, degron formation can explain some pathogenic gene variants. However, these are limited to regions that are exposed such as highly solvent accessible loops and intrinsically disordered regions.

## Discussion

Systematic mapping of degrons in proteins is essential for furthering our understanding of protein stability and abundance in the cellular environment, but as we show here, it may also provide mechanistic insight into how certain missense protein variants are pathogenic. We have comprehensively mapped degrons in all the human cytosolic proteins, thus providing a near complete map of degrons for 5,672 proteins corresponding to >25% of the protein coding genome. Our data support the idea that cells have a general quality control degradation mechanism, which mostly depends on the amino acid composition (7, 11, 15, 16, 31). This mechanism mainly recognizes hydrophobic residues, which normally are buried in the native protein structure. Several chaperones and E3s like HSP70, BAG6 and RNF126 (7, 17, 49) are potential candidates involved in this mechanism as they are likely to possess broad binding specificity. However, this composition-dependent degradation could also be an averaged effect of multiple different and more specific pathways, implicating a large variety of different chaperones, co-chaperones and ubiquitin ligases. Apart from the previously described C-degrons, we also noted some additional C-degrons and found C-terminal stabilization signals, including lysine, glutamine and acidic residues. Polyglutamine is known to be a difficult target for the proteasome (50) and found at the C-terminus of proteasome subunits to prevent their degradation by the proteasome (51). Similarly, an aspartate- and lysine-rich stabilizing peptide, derived from the C-terminus of the *Drosophila* proteasome subunit RPN10, has been shown to greatly stabilize proteins to which it was fused (52). The mechanistic details on how these C-terminal regions provide protein stabilization are unclear but they could in principle recruit deubiquitylating enzymes, shield against E3 binding, or provide poor degradation initiation sites for substrate unfolding by the proteasome.

We found that the average degron score correlates negatively with exposure and disorder in the native protein context. This agrees with previous observations, showing that PQC degrons are mostly buried within the structured regions of proteins (18, 42, 46) and thus only become relevant when they are exposed, e.g. due to misfolding. This also rationalizes our observation that the degron scores only correlate with full-length protein abundance when the degrons are found in highly exposed regions. We observed rapid proteasomal degradation of the keratin associated protein KRTAP11-1 that contains multiple exposed degrons. Presumably these degrons will be buried when the KRTAP11-1 interacts with keratin bundles, but these degrons likely also ensure that any unassembled KRTAP11-1 is degraded, as has been observed before (53).

In recent years there have been significant efforts towards identifying and characterizing degrons by both computational methods (16, 28, 29, 37, 54) and high-throughput experimental assays (7, 10, 11, 14, 15, 17, 30, 55–57). However, the relevance of the degron positioning in terms of exposure and structure in their native context has not been extensively studied. We used PAP to find protein regions where the abundance of single amino acid substitution variants can be explained by PQC degron creation or destruction and found that in exposed and unstructured regions the degron potency could largely explain the abundance of the full-length protein variants.

Thus, in the context of full-length PRKN, the effects of variants in the disordered linker region between the N-terminal UBL domain and the RING0 domain were efficiently captured by ΔPAP predictions, as were variants in most exposed loops in PRKN and ASPA. Therefore, if pathogenic PRKN and ASPA missense variants were found in these regions, they would likely result in insufficient protein levels, like our observations on the PNPO D33V variant. Since the D33V variant displays unchanged *k*_cat_ and thermal stability *in vitro* (48), the degron formation of the D33V substitution is likely the cause of the pathogenicity. Our analysis revealed several other possible candidates, where the mechanism of pathogenicity could involve the generation of a degron. These include HSPB1 T180I (ΔPAP = -0.086), a missense mutation that is associated with recessive inheritance of Charcot-Marie-Tooth disease and studies have revealed an increased thermal stability and no change of its chaperone-like activity (58). Another example is RARS1 D2G (ΔPAP = -0.077), which is associated with an autosomal recessive leukodystrophy and has been found to lead to defective assembly of the multi-synthetase complex, which could be due to a reduced expression (59). The fact that these diseases are recessive is also consistent with disease variants potentially generating degrons. These findings indicate that in combination with other variant effect predictors (VEPs), PAP may offer an orthogonal approach to clinical classification of variants observed in population sequencing, and importantly for exposed and disordered regions that are more poorly predicted by VEPs (60, 61).

However, not all variants in exposed and unstructured regions are captured by ΔPAP. As we found in ASPA, the correlation can break due to the presence of prolines, that upon removal would likely lead to structural destabilization of the protein. Moreover, in the case of PTEN, degron forming variants in its highly exposed and disordered C-terminal tail do not result in degradation of the full-length protein. This suggests that this region is either somehow inaccessible, e.g. through its known intramolecular interactions (62–64) or that its degron activity is reduced, e.g. by phosphorylation, where the negatively charged phosphate groups are likely to inhibit PQC degron activity. Indeed, PTEN is highly phosphorylated in the C-terminal region, and phosphorylation has been shown to block PTEN degradation (65). Thus, PAP can serve as a useful tool in the interpretation of high-throughput protein variant abundance assays.

The success of the proteolysis-targeting chimera (PROTAC) strategy for selective degradation of clinically relevant proteins (66), has recently led to a surge in interest in degrons. In general, PROTACs are heterobifunctional molecules, where one part of the molecule specifically interacts with the protein target, while the other part recruits an E3 ubiquitin-protein ligase, resulting in specific degradation of the target protein. Thus, furthering our understanding of degrons and E3 binding may increase the arsenal of E3s for developing PROTAC-based pharmaceuticals. As evident from the systematic screen that we present here, as well as from previous studies on large-scale degron mapping (7, 10, 11, 14, 15, 17, 30, 55, 57), the principal PQC degron signal is exposed hydrophobicity. Thus, a small molecule designed to specifically equip a target protein with a hydrophobic signal for PQC degradation could in principle achieve a similar outcome as a conventional PROTAC (66), and may hold the advantage of the apparent promiscuity of PQC E3s in terms of positioning of the substrate lysine residues for ubiquitylation.

## Methods

### Cell maintenance

The HEK293T TetBxb1BFPiCasp9 Clone 12 cell line (35) was grown in Dulbecco’s Modified Eagle’s Medium (DMEM) (Sigma-Aldrich) with 10% (v/v) fetal bovine serum (FBS) (Sigma Aldrich), 0.24 mg/mL streptomycin sulphate (BioChemica), 0.29 mg/mL penicillin G potassium salt (BioChemica), 0.32 mg/mL L-glutamine (Sigma Aldrich) and 2 µg/mL doxycycline (Dox) (Sigma-Aldrich). Cells were passaged when they reached 70-80% confluency (every two to three days). The cells were tested negative for mycoplasma (Mycostrip, InvivoGen). The authenticity of the cell line was ensured by regular selection of recombinant cells with 10 nM of AP1903 (MedChemExpress) and based on expression of BFP in the non-recombinant cells.

### Design of cytosolic library

Wild-type protein coding DNA sequences were downloaded from Ensembl (GRCh38.p13) via biomart on Sep. 15, 2020 (http://mart.ensembl.org/biomart/martview). Genes of cytosolic proteins were selected based on gene ontology annotation GO:0005829 (33). Sequences shorter than 90 nucleotides were discarded. For each of these 5672 genes, a transcript was selected based on available annotations from APPRIS, transcript support level (TSL) and GenCode (67, 68); see library/build_lib.r available on GitHub. Protein coding DNA sequences were sliced into tiles of 90 nucleotides with an overlap of 45. Substrate sequences of the NotI and BsiWI restriction enzymes were mutated to an alternative common codon as detailed in the supplementary material (**Table S1**). To avoid unintended template switching during amplification of DNA fragments, we split the tiles into 3 non-overlapping pools consisting of even, odd and C-terminal tiles. The even and odd libraries were further split randomly into two sub-libraries to get a manageable complexity. In addition to the 90-nt long tiles, five synonymous variants of five control tile sequences (25 control tiles in total) ranging from 66-up to 72-nt long tiles were also included in all the sublibraries to help compare scores across sublibraries. The control tiles were selected based on their scores from a previous assay we performed (42, 46). In total, we made five libraries consisting of Evens 1: 55184, Evens 2: 50270, Odds 3: 56582 Odds4: 51511 and C-terminal 1: 5223 unique DNA tiles. Due to isoforms and common domains, there is a small overlap between pairs of tile libraries and a few synonymous sequences within libraries. The 5672 protein-coding genes from Ensembl (5321 unique; out of ∼70,000 human protein coding genes) were mapped to 5129 non-redundant proteins from the UniProt human proteome (out of ∼23,000) and thus represents 22% of the human proteome.

### Cytosolic library cloning

The five libraries, containing a total of 218,770 tiles (213,261 unique), were purchased from Twist Bioscience. The oligos were designed with two additional adaptors that included complementary sequences (5’ complementary sequence: GTTCTAGAGGCAGCGGAGCCACC, 3’ complementary sequence: TAGTAACTTAAGAATTCACCGGTCTGACCT) to the flanking region of the ‘attB-EGFP-PTEN-IRES-mCherry-562bgl’ (34) (p2127) vector, where they were inserted. A NotI (GCGGCCGC) or a BsiWI (CGTACG) restriction site was included upstream and downstream of these complementary sequences respectively. Upstream of the NotI restriction site and downstream of the BsiWI restriction site specific primer binding sites (see primers VV7S-VV14S in **Supplementary Data File**) were designed to allow for independent amplification of two sublibraries from the Evens and the Odds and the amplification of the one C-terminal library. The index number of each library is indicative of the primer set used to amplify them.

Each sublibrary of oligos were amplified by PCR using Q5 High-Fidelity 2x Mastermix (New England Biolabs) with the primers VV7S-VV14S by denaturing at 98 °C for 30 s; followed by cycling for 12 times at 98 °C for 5 s, 59 °C for 30 s, 72 °C for 10 s, with a final extension step at 72 °C for 5 min. The p2127 plasmid was amplified and linearized by PCR with Q5 High-Fidelity 2x Mastermix by the primers VV1 and VV2 with the following PCR program: 98 °C for 30 s; followed by cycling for 30 times at 98 °C for 5 s, 69 °C for 30 s, 72 °C for 3 min and 40 s, with a final extension step at 72 °C 5 min. Once amplified, both oligos and the vector were purified by Zymo Clean and Concentrator-5 kit (Zymo Research) and eluted in nuclease-free water (Thermo Fisher Scientific) (30 μL for the amplified linear vector and 10 μL for the amplified oligos). To eliminate the PCR template for the vector PCR, the purified PCR was digested with DpnI (New England Biolabs). The amplified sublibraries of oligos were digested by NotI (New England Biolabs) and BsiWI (New England Biolabs). The digested vector PCR and the digested oligo PCRs were run on an agarose gel (1.5% for vector PCR and 2% for oligo PCRs). DNA was stained with 1x SYBR Safe (Thermo Fisher Scientific). The properly sized DNA bands were extracted from the agarose gels with GeneJET Gel Extraction Kit (Thermo Scientific). The DNA concentration of the extracted bands was measured by Qubit 1X dsDNA High Sensitivity (HS) Kit (Thermo Fisher Scientific).

The linearized p2127 vector was mixed with each of the amplified oligo sublibraries with a molecular ratio of 1:4 to perform a Gibson assembly reaction (New England Biolabs) following the manufacturer’s instructions. The Gibson assembly reaction was purified with the Zymo Clean and Concentrator-5 kit and eluted in 12 μL of nuclease-free water.

NEB-10β *E. coli* cells (New England Biolabs) (50 μL) were transformed by electroporation at 2 kV with 2 μL of the cleaned Gibson assembly reaction, according to the manufacturer’s protocol. Then, 11 transformations per Evens and Odds sublibrary were performed along with 5 transformations for the C-terminals, to ensure complete coverage of the sublibraries. A small portion (1/11,000 or 1/5,000 for the C-terminals) of the transformed cells were plated on LB-agar plates containing ampicillin (100 μg/mL) and left to grow at 37 °C overnight. At the same time the rest of the cells were inoculated into 200 mL of LB media with ampicillin and cultured overnight at 37 °C. The next day, the colonies were counted to ensure at least 100-fold coverage of the complexity of each sublibrary. Midipreps were performed on the 200 mL cultures with the NucleoBond Xtra Midi kit (Macherey-Nagel). The DNA yield and purity from the midiprep cultures was measured by the NanoDrop spectrometer ND-1000.

### Cloning of individual tile and protein constructs

All tiles that were individually assessed were synthesized by IDT, with the exception of the tiles for the ASPA protein (from Eurofins). Then they were cloned into p2127 with the same Gibson assembly cloning strategy that was used for the library cloning. All the full-length proteins (same isoforms that were used for the library construction) were synthesized and cloned into the p2127 vector by Genscript. The tiles used for verification of the abundance scores were picked randomly from the transformation plates of the five sublibraries. The PPARG and NR3C1 protein tiles were synthesized and cloned by Genscript.

### Transfections

Transfections of the sublibraries and of different plasmids encoding individual tiles or protein variants were performed in 15 cm plates (8 x 10^6^ cells) (for sublibraries) or 12-well plates (0.2 x 10^6^ cells), or 96-wel plates (20 x 10^3^ cells) (for individual construct transfections) in cell cultures of HEK293T TetBxb1BFPiCasp9 Clone 12 cells without Dox. The library or individual tile/variant plasmid was mixed with the pCAG-NLS-Bxb1 (Addgene, Plasmid #51271) plasmid at a molar ratio of 17.5:1 along with OptiMEM (Thermo Fisher Scientific) (3,570 μL for 15 cm plate and proportionately less for the other plates based on their surface area) and Fugene HD (Promega) (73 μL for a 15 cm plate). The mixture was incubated for 15 min and added to the cells. After 48 h Dox (2 μg/mL) was reintroduced in the cell culture, as well as 10 nM of AP1903 to ensure counterselection of non-recombinant cells. The cells for sublibrary transfections were grown for another 4 days, while passaging after two days to allow for their tiles to reach a steady state level. For the individual transfections the number of days that the cells were allowed to grow after reintroduction of Dox and AP1903 treatment varied between 4 and 10 days.

### Cell perturbations

The Evens 1 sublibrary was used to evaluate the broad effects of different drug treatments on the library tiles. The sublibrary was treated with 10 μg/mL of cycloheximide for 8 hours (Sigma Aldrich). It was treated with 15 μM of Bortezomib (BZ) (LC Laboratories) for 16 hours and with 20 μM of chloroquine (CQ) (Sigma Aldrich). Finally, the sublibrary was treated with 1 μM of MLN7243 (MedChemExpress) and 1.5 μM of MLN4924 (MedChemExpress) for 16 h. Addition of the equivalent volume of DMSO, which is the solvent of BZ, MLN7243 and MLN4924 showed no effect in comparison to not adding anything in the culture.

### Flow cytometry and flow sorting

After transfection and allowing the expressed tile/variant to reach steady state levels in the cell, flow cytometry was performed using either a BD FACSJazz flow cell sorter (for 12-well plates) or the BD LSRFortessa cell analyzer (for 96-well plates).

For the 15 cm plate transfections of the sublibraries, experiments were performed in three biological replicates for each sublibrary and two sorting replicates per biological replicate (in total six replicates per sublibrary). The sorting of the cells was performed with the BD FACSAria III cell sorter and cells were sorted into four equally populated bins. The cells were prepared by washing with PBS, trypsinizing, centrifuging (300 g, 5 min) and then resuspending cells in PBS with 2% FBS. Cells were sorted in tubes with 1 mL of cell growth medium. After sorting, the cells were precipitated and resuspended in medium without Dox where they were allowed to grow for 4 to 6 days. Subsequently, the cells were harvested in 50 mL tubes and stored at -80 °C for future genomic DNA extraction. To ensure full coverage of the sublibraries at least 100x of the complexity of each sublibrary of cells were maintained at each stage.

For excitation of GFP, mCherry and BFP 488 nm, 561 nm and 405 nm lasers were used respectively. For BD FACSJazz the filters used were 530/40, 610/20 and 450/50 respectively. For BD LSRFortessa the corresponding filters were 530/30, 610/20 and 431/28. Finally, for BD FACSAriaIII the equivalent filters were 530/30, 615/20 and 442/46. The backgating for both sorting and analyzing cells involved gating based on FSC and SSC for live cells, gating based on the FSC width for singlets and gating the mCherry positive and BFP negative for selection of the recombinant cells. Examples of the gating strategies used are included in the supplementary material (**Fig. S18**).

### Illumina sequencing of sorted cells

For each sorted bin of each replicate, genomic DNA was extracted from approximately 30 x 10^6^ cells with the DNeasy blood & tissue Kit (Qiagen). A fraction of the extracted DNA was used to perform 16 PCRs (2 PCRs for the C-terminals) using 2.5 μg of DNA per 50 μL PCR as template for each bin. The PCRs were performed using Q5 High-Fidelity 2x Mastermix (New England Biolabs) and the primers VV7 and VV8. The PCR program consisted of initial denaturation at 98 °C for 30 s; followed by cycling for 7 cycles at 98 °C for 5 s, 65 °C for 30 s, 72 °C for 50 s, with a final extension step at 72 °C, 2 min. The PCR product was purified by Ampure XP beads (Beckman Coulter) (0.8:1 volume ratio of PCR to reagent) and eluted in 20 μL. A second PCR was performed using as template 4 μL of the purified first PCR product, in order to add to it demultiplexing indices and the cluster generating sequences for subsequent Illumina sequencing. The PCRs were performed using Q5 High-Fidelity 2x Mastermix (New England Biolabs) and one of the primers VV29-VV31 as forward and one of JS1-JS50 as reverse. The PCR program was an initial denaturation at 98 °C for 30 s; followed by cycling for 14 cycles at 98 °C for 5 s, 67 °C for 30 s and 72 °C for 15 s. The amplicons were applied to a 2% agarose gel with 1x SYBR Safe (Thermo Fisher Scientific) and extracted using the GeneJET gel extraction kit (Themo Scientific). The purified DNA bands were quantified by Qubit 2.0 Fluorometer (Invitrogen), using the Qubit dsDNA HS Assay Kit (Thermo Fisher Scientific). Taking into account the complexity of each sublibrary, the prepared amplicons of each bin were mixed so that at least 100x coverage of their equivalent sublibrary complexity was ensured. The mixed solution of all the amplicons was loaded on a NextSeq 500/550 High output (300 cyles) kit (Illumina), along with 10% PhiX (Illumina) and they were sequenced with a 550 NextSeq 550 system (Illumina). Custom Read and Index primers were used for the sequencing runs: Read 1: VV17, Read2: VV25, Index 1: VV19 and Index 2: VV16.

### Illumina sequencing processing and degron score calculation

Demultiplexing of the sequenced tiles was performed with the BaseSpace Sequence Hub. Adapter sequences were removed using cutadapt (69) and paired-end reads were joined using fastq-join from ea-utils (70), see counts/call_zerotol_paired.sh available at GitHub. Only sequences with a perfect match were counted, see scores/merge_and_map.r. Read counts from 120 pairs of technical replicates have an average Pearson correlation of 0.98 (range 0.79-0.99) and an average of 3.8 mill. matched reads per technical replicate. Technical replicates were merged and normalized to frequencies without pseudo counts. For each biological and FACS replicates, we calculated a protein stability index (PSI) per tile, *t*, using:

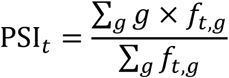

where *f_t,g_* is the frequency of tile *t* in FACS gate *g*, and requiring a total of 100 or more reads across the four gates. All individual replicates had scores calculated for >99% of library members. Each of the five sub-libraries have 6 replicates, 3 biological times 2 FACS replicates, with an average PSI Pearson correlation of 0.98 between replicate pairs (range 0.96-0.99, average over 15 pairs times 5 sub-libraries).

Before merging PSI scores from different libraries and replicates, it is important to ensure that these are comparable scores. The PSI scale is related to the FACS gating which is set to fill 25% of cells in each of the four bins. Because most FACS runs are observed to have similar overall fluorescence distribution, the gating should be similar in each sorting experiment and thus, the PSI scores should be close to comparable. Each replicate is normalized to account for minor variations by aligning PSI distributions using the first and last peak located as the median value of the first and last quarter of the scores. In total, 4,573 pairs of identical tiles measured in two different libraries were available and the Pearson correlation between PSI and peak normalized scores were all >0.99. To merge replicates and libraries, normalized degron scores were averaged per tile and the standard deviation between replicates reported as uncertainty (mean 0.05, range 0.00-0.33). We require two or more replicate measurements per tile resulting in 212,658 scores and a final coverage of 99.7%.

### Transformation of degron score distribution

As discussed above, we consider the degron score distribution with four peaks a consequence of the sort-seq experiment. To get a more physically relevant scale, we transformed the degron scores such that their distribution matches the fluorescence distribution from flow cytometry while maintaining the same order. The resulting abundance score has reverse ordering compared to the degron score (Spearman correlation -1) and is used for training models such that these predict abundance on a scale proportional to the GFP/mCherry fluorescence ratio. To carry out this transformation, we first make quantile function of the flow cytometry (FC) distribution implemented as a spline function of the inverse cumulated density. To avoid boundary effects, we add 2% of a normal distribution with the same mean and standard deviation as the FC distribution. The quantile function may be used to transform uniformly distributed numbers to the FC distribution. The degron scores can also be transformed to a uniform distribution using the cumulated density, also implemented as a spline function, and together these transform the degron score distribution to the FC distribution, see score/scores.r available on GitHub. The derivative of the combined function is used to propagate uncertainties.

### Holdout test data

Before doing any modeling, we carefully select a fraction of the data for a final validation. Full protein sequences were clustered using MMseqs2 (71) and 527 proteins (∼10%) were selected at random among non-redundant proteins (cluster size one). Of these, 99 proteins were re-assigned as training data because they had tiles that were similar to other training tiles resulting in 7% of tiles being assigned as holdout test data. See library/build_lib.r available on GitHub. In the final set of 211,444 measured tiles with unique amino acid sequences used for models, the test data constitutes 15,815 tiles (7.5%), see models/model_data.r.

### Logistic regression model

We have previously found that a logistic regression model using amino acid composition is robust even when the true distribution of the data is not known (16). For comparison, we make a similar model here by considering only the composition of internal region of the peptides thus avoiding C-degron effects. A binary label is made by splitting the data at the median score and fitting a model to the amino acid composition of position 1-25, see models/regression_analysis.r available on GitHub. The predicted degron probability from this model is observed to correlate well with degron score (Pearson -0.81) and abundance score (Pearson 0.82 and Spearman 0.82) on the test data and despite being a classifier response without a C-degron term. The model is included in the PAP software under the name “human25”.

### Linear regression analysis

We carried out a two-step linear regression analysis to determine both the importance and the strength of sequence features. First, we used lasso regression to order sequence features according to predictive power by scanning the regularization strength from 10^-1^ to 10^-6^ in 501 logarithmically spaced steps. Features with non-zero coefficients at higher regularization strength were considered more important. Second, selections of features were used in non-regularized linear regressions from which the coefficients were considered the strength of the feature. The statistical significance of features was determined based on the non-regularized regression using *t*-statistics and corresponding two-sided *p*-value. This analysis was carried out using the *R project for statistical computing* with the package *glmnet*.

We systematically tested many sequence features in a single regression. Twenty composition features were the summed number of each amino acid in each peptide. All position-specific effects of each amino acid were included with 600 features. A peptide of 30 residues has 174,000 position specific pairs. To reduce this number to a more manageable set of residue pairs we first calculated the score coupling, *c_a1,a2_*, of all pairs using:

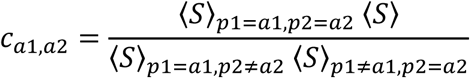

where 〈*S*〉 is the score average for the specified peptides, e.g. position *p1* being amino acid *a1*. Couplings were only calculated if all averages were over 100 or more peptides. *c*_a1,a2_ thus helps us select amino acid/position pairs where the abundance effect is less well described by the individual amino acids and positions. From this we selected the pairs with more than 20% coupling, i.e. features with coupling greater than 1.2 or smaller than 0.8, resulting in 522 destabilizing and 1154 stabilizing pair features. Similarly, for position specific triplets we considered the last 7 positions and we calculated 280,000 “ternary” couplings using:

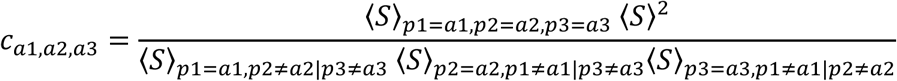

where e.g. *p*1 = *a*1, *p*2 ≠ *a*2|*p*3 ≠ *a*3 denotes the peptides that have amino acid *a1* at position *p1* and possibly one more amino acid of the triplet at its respective position, but not all three. From this we selected 212 destabilizing and 180 stabilizing triplet features, again using a threshold of 100 occurrences and 20% coupling. For both pairs and triplets, the calculated couplings are only used to reduce the number of features in the lasso analysis. In addition to the lasso regression with all features, we constructed two linear models with selected features and without regularization. One model included the maximum number, 179, of top-ranking significant features (p-value < 0.01 for all coefficients) resulting in a Pearson correlation of 0.81 (**Fig. S11**). The other model included a smaller selection of 55 features but similar correlation (Pearson 0.80 and Spearman 0.85) and is available in the PAP software under the name “human30”. This model only contained position specific features in the 5 last C-terminal positions, and we consider these a C-degron term and the composition of the full peptide an internal term. Using both terms correlated better with abundance score (Pearson 0.80 and Spearman 0.86) compared to the internal term alone (Pearson 0.76 and Spearman 0.82) when applied to the holdout test data. Features and coefficients of all models are available in supplementary material (**Supplementary Data File**).

### Two-way convolutional neural network

Based on the regression analysis we aimed to make a model that more accurately captures the score distribution and non-linear effects. Furthermore, we aimed for a model with the ability to evaluate peptides with and without C-degron effects and peptides of slightly different lengths. Hypothesizing that internal degradation signals are local in sequences and translationally invariant we use a convolutional neural network (CNN) filter that strides over the peptide with a padding of one and globally pooled to a single set of filters. C-degron effects, on the other hand, are localized to the C-terminal positions and thus we add a second channel that only considers these. The two channels are followed by a number of dense layers and each summed to a single number that describes the internal effect and C-degron effect. Finally, the outputs of the two channels are added to give the observed score (**Fig. 4A**). In the regression model, 27% (77%) of peptides have a C-degron score <0.01 (0.05) and these should ensure that the internal channel is well calibrated. We used 20% peptides for validation (different from the holdout test data described above) and early stopping with a maximum of 2000 epochs.

For optimizing hyperparameters, we first scanned various hyperparameters with 1–4 dense layers following the input filters in each channel. No improvement in mean square error (MSE) loss was observed with more than two dense layers. Using training parameters found to be robust in the first scans, we next scanned hyperparameters related to the input filters; the number (25, 50, 100, 150) and size (1–21 residues) of CNN filters; the number (25, 50, 100, 150) and size (5–7 residues) of C-degron filters. From the regression analysis, we expected C-degron effects to be mainly destabilizing but for most architectures we observed a positive median score from the C-degron channel. The sign of the median C-degron score varied although seemingly influenced by some hyperparameters. Thus, we used the following three-step training scheme: First, the CNN channel was trained on positions 2-25 only (the central positions with similar behavior) and without a C-degron channel. Second, the C-degron channel was trained while keeping the CNN channel fixed but now given all positions as input. Third, the dense layers of both channels were refined with a lower learning rate. Based on this training scheme, we selected 25 CNN filters of one residue and 25 C-degron filters of 5 residues. Finally, training parameters were scanned; learning rate (1e-3, <u>1e-4</u>, 1e-5); dropout rate (0.0, 0.1, <u>0.2</u>); l2 regularization weight (1e-5, <u>1e-6</u>, 1e-7); the number of filters in dense layers (50, 100, 150); batch size (512, 1024, 2048) and activation function (<u>ELU</u>, sigmoid). The selected values are underlined. This is the default model in the PAP software and named “cnn2w1”.

### Structural analyses

Protein structures were obtained from version 4 of the AlphaFold2 Protein Structure Database (AFDBv4) (https://ftp.ebi.ac.uk/pub/databases/alphafold/v4/) (72, 73). Proteins with fewer than 16 residues, those containing non-standard amino acids, or those absent from the AlphaFold UniProt reference proteome “one sequence per gene” FASTA file were excluded. For proteins longer than 2,700 amino acids, whose AFDBv4 predictions are split into 1,400-residue segments with a 1,200-residue overlap, we calculated exposure and extracted pLDDT values for each segment, then averaged the overlapping residues at each position to obtain a single representative value. We retrieved the predicted local distance difference test (pLDDT) scores from AlphaFold2 (AF2) structures, using these scores as a proxy for local structural confidence and possible disorder (74). Based on the AF2 models, we calculated a descriptor of residue exposure by determining the relative Accessible Surface Area (rASA) using DSSP via BioPython v1.81 with default settings (75).

For the structural analysis of degron scores we made a “cytosolic proteome” dataset consisting of all tiles from 5100 proteins for which we were able to map a AF2 structure. Degrons scores were copied to all occurrences of redundant tiles. Average rASA and pLDDT values of all residues in a tile were calculated to represent tiles.

### ClinVar analysis

ClinVar data were obtained from the publicly available variant summary file (e.g., variant_summary_2024-03.txt.gz) at ftp://ftp.ncbi.nlm.nih.gov/pub/clinvar/tab_delimited/ on [2024-03-18]. Each full-length protein sequence in our library was aligned to its corresponding ClinVar reference sequence, and only proteins with sequences identical to the ClinVar reference were retained. Entries with a review status below one star, insertions or deletions, or mutations altering start codons were excluded. Additionally, only those variants annotated as “benign,” “likely benign,” “pathogenic,” or “likely pathogenic” were kept, while all other clinical significance categories were removed. To harmonize the final labels, “benign” variants were merged with those labelled as “likely benign,” and “pathogenic” variants were merged with the “likely pathogenic” variants. PAP scores of WT and mutant were calculated using a 30-residue window around the mutation site without the C-degron term (unless the tile ends at the C-terminus). The resulting dataset is provided in the supplementary data file. For analysis of residue exposure in clinical variants, the average rASA of the variant itself and the five residues on either side were used to derive a single exposure value.

## Data availability

All data and software generated for this article is available on GitHub: https://github.com/KULL-Centre/_2025_Voutsinos_degron_cytosol. Sequencing reads, counts and FACS files are available at https://sid.erda.dk/sharelink/FUolgBZn9W. An online accessible version of PAP is available via Google Colab (see GitHub link above).

## Acknowledgements

We acknowledge the use of the FACS, sequencing and computing core facilities at the Biotech Research & Innovation Centre and Department of Biology, University of Copenhagen. We thank Vibe H. Oestergaard, Michael Lisby, Søren Lindemose, Danai Trimintziou and Anne-Marie Lauridsen for technical assistance.

## Supplementary material

This article includes the following supplementary information:

- Supplementary Figures and Tables
- Source Data (SupplementaryDataFile.xlsx)

## Author contributions

V.V., K.E.J., F.B.L., M.G.-T., N.J., E.H.-O., performed the experiments. V.V., K.E.J., N.J., G.T., A.S., D.M.F., K.L.-L., and R.H.-P. analyzed the data. K.L.-L. and R.H.-P. conceived the study. V.V., K.E.J., and R.H.-P. wrote the paper.

## References

1. Hershko A, Ciechanover A. The ubiquitin system. Annu Rev Biochem. 1998;67:425–79.

2. Bard JAM, Goodall EA, Greene ER, Jonsson E, Dong KC, Martin A. Structure and Function of the 26S Proteasome. Annu Rev Biochem. 2018;87:697–724.

3. Schimke RT, Doyle D. Control of enzyme levels in animal tissues. Annu Rev Biochem. 1970;39:929–76.

4. Mathieson T, Franken H, Kosinski J, Kurzawa N, Zinn N, Sweetman G, et al. Systematic analysis of protein turnover in primary cells. Nat Commun. 2018;9(1):689.

5. Eden E, Geva-Zatorsky N, Issaeva I, Cohen A, Dekel E, Danon T, et al. Proteome half-life dynamics in living human cells. Science. 2011;331(6018):764–8.

6. Ravid T, Hochstrasser M. Diversity of degradation signals in the ubiquitin-proteasome system. Nature reviews Molecular cell biology. 2008;9(9):679–89.

7. Zhang Z, Sie B, Chang A, Leng Y, Nardone C, Timms RT, et al. Elucidation of E3 ubiquitin ligase specificity through proteome-wide internal degron mapping. Mol Cell. 2023;83(18):3377–92.e6.

8. Hartooni N, Sung J, Jain A, Morgan DO. Single-molecule analysis of specificity and multivalency in binding of short linear substrate motifs to the APC/C. Nat Commun. 2022;13(1):341.

9. Lee JM, Hammarén HM, Savitski MM, Baek SH. Control of protein stability by post-translational modifications. Nat Commun. 2023;14(1):201.

10. Timms RT, Zhang Z, Rhee DY, Harper JW, Koren I, Elledge SJ. A glycine-specific N-degron pathway mediates the quality control of protein N-myristoylation. Science. 2019;365(6448).

11. Koren I, Timms RT, Kula T, Xu Q, Li MZ, Elledge SJ. The Eukaryotic Proteome Is Shaped by E3 Ubiquitin Ligases Targeting C-Terminal Degrons. Cell. 2018;173(7):1622–35.e14.

12. Varshavsky A. N-degron and C-degron pathways of protein degradation. Proc Natl Acad Sci U S A. 2019;116(2):358–66.

13. Timms RT, Koren I. Tying up loose ends: the N-degron and C-degron pathways of protein degradation. Biochemical Society Transactions. 2020;48(4):1557–67.

14. Maurer MJ, Spear ED, Yu AT, Lee EJ, Shahzad S, Michaelis S. Degradation signals for ubiquitin-proteasome dependent cytosolic protein quality control (CytoQC) in yeast. G3: Genes, Genomes, Genetics. 2016;6(7):1853–66.

15. Mashahreh B, Armony S, Johansson KE, Chappleboim A, Friedman N, Gardner RG, et al. Conserved degronome features governing quality control associated proteolysis. Nat Commun. 2022;13(1):7588.

16. Johansson KE, Mashahreh B, Hartmann-Petersen R, Ravid T, Lindorff-Larsen K. Prediction of Quality-control Degradation Signals in Yeast Proteins. J Mol Biol. 2023;435(2):167915.

17. Abildgaard AB, Voutsinos V, Petersen SD, Larsen FB, Kampmeyer C, Johansson KE, et al. HSP70-binding motifs function as protein quality control degrons. Cellular and molecular life sciences : CMLS. 2023;80(1).

18. Kampmeyer C, Larsen-Ledet S, Wagnkilde MR, Michelsen M, Iversen HKM, Nielsen SV, et al. Disease-linked mutations cause exposure of a protein quality control degron. Structure (London, England : 1993). 2022;30(9):1245-53.e5.

19. Stein A, Fowler DM, Hartmann-Petersen R, Lindorff-Larsen K. Biophysical and Mechanistic Models for Disease-Causing Protein Variants. Trends in biochemical sciences. 2019;44(7):575–88.

20. Thrun A, Garzia A, Kigoshi-Tansho Y, Patil PR, Umbaugh CS, Dallinger T, et al. Convergence of mammalian RQC and C-end rule proteolytic pathways via alanine tailing. Mol Cell. 2021;81(10):2112–22.e7.

21. Shen PS, Park J, Qin Y, Li X, Parsawar K, Larson MH, et al. Protein synthesis. Rqc2p and 60S ribosomal subunits mediate mRNA-independent elongation of nascent chains. Science. 2015;347(6217):75-8.

22. Howard CJ, Frost A. Ribosome-associated quality control and CAT tailing. Crit Rev Biochem Mol Biol. 2021;56(6):603–20.

23. Kostova KK, Hickey KL, Osuna BA, Hussmann JA, Frost A, Weinberg DE, et al. CAT-tailing as a fail-safe mechanism for efficient degradation of stalled nascent polypeptides. Science. 2017;357(6349):414–7.

24. Sitron CS, Brandman O. CAT tails drive degradation of stalled polypeptides on and off the ribosome. Nat Struct Mol Biol. 2019;26(6):450–9.

25. Patil PR, Burroughs AM, Misra M, Cerullo F, Insua CC, Hung HC, et al. Mechanism and evolutionary origins of alanine-tail C-degron recognition by E3 ligases Pirh2 and CRL2-KLHDC10. Cell Rep. 2023;42(9):113100.

26. Liu Y, Chen J, Khusnutdinova AN, Correia K, Diep P, Batyrova KA, et al. A novel C-terminal degron identified in bacterial aldehyde decarbonylases using directed evolution. Biotechnol Biofuels. 2020;13:114.

27. Zutz A, Hamborg L, Pedersen LE, Kassem MM, Papaleo E, Koza A, et al. A dual-reporter system for investigating and optimizing protein translation and folding in E. coli. Nat Commun. 2021;12(1):6093.

28. Tokheim C, Wang X, Timms RT, Zhang B, Mena EL, Wang B, et al. Systematic characterization of mutations altering protein degradation in human cancers. Mol Cell. 2021;81(6):1292–308.e11.

29. Hou C, Li Y, Wang M, Wu H, Li T. Systematic prediction of degrons and E3 ubiquitin ligase binding via deep learning. BMC Biol. 2022;20(1):162.

30. Geffen Y, Appleboim A, Gardner RG, Friedman N, Sadeh R, Ravid T. Mapping the Landscape of a Eukaryotic Degronome. Mol Cell. 2016;63(6):1055–65.

31. Timms RT, Mena EL, Leng Y, Li MZ, Tchasovnikarova IA, Koren I, et al. Defining E3 ligase-substrate relationships through multiplex CRISPR screening. Nat Cell Biol. 2023;25(10):1535–45.

32. Makaros Y, Raiff A, Timms RT, Wagh AR, Gueta MI, Bekturova A, et al. Ubiquitin-independent proteasomal degradation driven by C-degron pathways. Mol Cell. 2023;83(11):1921–35.e7.

33. Ashburner M, Ball CA, Blake JA, Botstein D, Butler H, Cherry JM, et al. Gene ontology: tool for the unification of biology. The Gene Ontology Consortium. Nat Genet. 2000;25(1):25–9.

34. Matreyek KA, Starita LM, Stephany JJ, Martin B, Chiasson MA, Gray VE, et al. Multiplex assessment of protein variant abundance by massively parallel sequencing. Nature genetics. 2018;50(6):874–82.

35. Matreyek KA, Stephany JJ, Chiasson MA, Hasle N, Fowler DM. An improved platform for functional assessment of large protein libraries in mammalian cells. Nucleic acids research. 2020;48(1).

36. Gersing SK, Wang Y, Grønbæk-Thygesen M, Kampmeyer C, Clausen L, Willemoës M, et al. Mapping the degradation pathway of a disease-linked aspartoacylase variant. PLoS Genet. 2021;17(4):e1009539.

37. Szulc NA, Stefaniak F, Piechota M, Soszyńska A, Piórkowska G, Cappannini A, et al. DEGRONOPEDIA: a web server for proteome-wide inspection of degrons. Nucleic Acids Res. 2024.

38. Kussie PH, Gorina S, Marechal V, Elenbaas B, Moreau J, Levine AJ, et al. Structure of the MDM2 oncoprotein bound to the p53 tumor suppressor transactivation domain. Science. 1996;274(5289):948–53.

39. Busino L, Donzelli M, Chiesa M, Guardavaccaro D, Ganoth D, Dorrello NV, et al. Degradation of Cdc25A by beta-TrCP during S phase and in response to DNA damage. Nature. 2003;426(6962):87–91.

40. Zhuang M, Calabrese MF, Liu J, Waddell MB, Nourse A, Hammel M, et al. Structures of SPOP-substrate complexes: insights into molecular architectures of BTB-Cul3 ubiquitin ligases. Mol Cell. 2009;36(1):39–50.

41. Zhao S, Olmayev-Yaakobov D, Ru W, Li S, Chen X, Zhang J, et al. Molecular basis for C-degron recognition by CRL2(APPBP2) ubiquitin ligase. Proc Natl Acad Sci U S A. 2023;120(43):e2308870120.

42. Grønbæk-Thygesen M, Voutsinos V, Johansson KE, Schulze TK, Cagiada M, Pedersen L, et al. Deep mutational scanning reveals a correlation between degradation and toxicity of thousands of aspartoacylase variants. Nat Commun. 2024;15(1):4026.

43. Boyle GE, Sitko K, Galloway JG, Haddox HK, Bianchi AH, Dixon A, et al. Deep mutational scanning of CYP2C19 reveals a substrate specificity-abundance tradeoff. bioRxiv. 2023.

44. Amorosi CJ, Chiasson MA, McDonald MG, Wong LH, Sitko KA, Boyle G, et al. Massively parallel characterization of CYP2C9 variant enzyme activity and abundance. Am J Hum Genet. 2021;108(9):1735–51.

45. Suiter CC, Moriyama T, Matreyek KA, Yang W, Scaletti ER, Nishii R, et al. Massively parallel variant characterization identifies NUDT15 alleles associated with thiopurine toxicity. Proc Natl Acad Sci U S A. 2020;117(10):5394–401.

46. Clausen L, Voutsinos V, Cagiada M, Johansson KE, Grønbæk-Thygesen M, Nariya S, et al. A mutational atlas for Parkin proteostasis. Nat Commun. 2024;15(1):1541.

47. Chiasson MA, Rollins NJ, Stephany JJ, Sitko KA, Matreyek KA, Verby M, et al. Multiplexed measurement of variant abundance and activity reveals VKOR topology, active site and human variant impact. Elife. 2020;9.

48. Barile A, Mills P, di Salvo ML, Graziani C, Bunik V, Clayton P, et al. Characterization of Novel Pathogenic Variants Causing Pyridox(am)ine 5’-Phosphate Oxidase-Dependent Epilepsy. Int J Mol Sci. 2021;22(21).

49. Müller MBD, Kasturi P, Jayaraj GG, Hartl FU. Mechanisms of readthrough mitigation reveal principles of GCN1-mediated translational quality control. Cell. 2023;186(15):3227–44.e20.

50. Venkatraman P, Wetzel R, Tanaka M, Nukina N, Goldberg AL. Eukaryotic proteasomes cannot digest polyglutamine sequences and release them during degradation of polyglutamine-containing proteins. Mol Cell. 2004;14(1):95–104.

51. Singh Gautam AK, Yu H, Yellman C, Elcock AH, Matouschek A. Design principles that protect the proteasome from self-destruction. Protein Sci. 2022;31(3):556–67.

52. Rethi-Nagy Z, Abraham E, Udvardy K, Klement E, Darula Z, Pal M, et al. STABILON, a Novel Sequence Motif That Enhances the Expression and Accumulation of Intracellular and Secreted Proteins. Int J Mol Sci. 2022;23(15).

53. Fujimoto S, Takase T, Kadono N, Maekubo K, Hirai Y. Krtap11-1, a hair keratin-associated protein, as a possible crucial element for the physical properties of hair shafts. J Dermatol Sci. 2014;74(1):39–47.

54. Martínez-Jiménez F, Muiños F, López-Arribillaga E, Lopez-Bigas N, Gonzalez-Perez A. Systematic analysis of alterations in the ubiquitin proteolysis system reveals its contribution to driver mutations in cancer. Nat Cancer. 2020;1(1):122–35.

55. Kats I, Khmelinskii A, Kschonsak M, Huber F, Knieß RA, Bartosik A, et al. Mapping Degradation Signals and Pathways in a Eukaryotic N-terminome. Mol Cell. 2018;70(3):488–501.e5.

56. Kong KE, Shankar S, Rühle F, Khmelinskii A. Orphan quality control by an SCF ubiquitin ligase directed to pervasive C-degrons. Nat Commun. 2023;14(1):8363.

57. Sharninghausen R, Hwang J, Dennison DD, Baldridge RD. Identification of ERAD-dependent degrons for the endoplasmic reticulum lumen. bioRxiv. 2024.

58. Chalova AS, Sudnitsyna MV, Strelkov SV, Gusev NB. Characterization of human small heat shock protein HspB1 that carries C-terminal domain mutations associated with hereditary motor neuron diseases. Biochim Biophys Acta. 2014;1844(12):2116–26.

59. Nafisinia M, Sobreira N, Riley L, Gold W, Uhlenberg B, Weiß C, et al. Mutations in RARS cause a hypomyelination disorder akin to Pelizaeus-Merzbacher disease. Eur J Hum Genet. 2017;25(10):1134–41.

60. Fawzy M, Marsh JA. Understanding the heterogeneous performance of variant effect predictors across human protein-coding genes. Sci Rep. 2024;14(1):26114.

61. Cagiada M, Jonsson N, Lindorff-Larsen K. Decoding molecular mechanisms for loss of function variants in the human proteome. bioRxiv. 2024.

62. Malaney P, Pathak RR, Xue B, Uversky VN, Davé V. Intrinsic disorder in PTEN and its interactome confers structural plasticity and functional versatility. Sci Rep. 2013;3:2035.

63. Rahdar M, Inoue T, Meyer T, Zhang J, Vazquez F, Devreotes PN. A phosphorylation-dependent intramolecular interaction regulates the membrane association and activity of the tumor suppressor PTEN. Proc Natl Acad Sci U S A. 2009;106(2):480–5.

64. Ross AH, Gericke A. Phosphorylation keeps PTEN phosphatase closed for business. Proc Natl Acad Sci U S A. 2009;106(5):1297–8.

65. Torres J, Pulido R. The tumor suppressor PTEN is phosphorylated by the protein kinase CK2 at its C terminus. Implications for PTEN stability to proteasome-mediated degradation. J Biol Chem. 2001;276(2):993–8.

66. Verma R, Mohl D, Deshaies RJ. Harnessing the Power of Proteolysis for Targeted Protein Inactivation. Mol Cell. 2020;77(3):446–60.

67. Rodriguez JM, Maietta P, Ezkurdia I, Pietrelli A, Wesselink JJ, Lopez G, et al. APPRIS: annotation of principal and alternative splice isoforms. Nucleic Acids Res. 2013;41(Database issue):D110–7.

68. Frankish A, Diekhans M, Ferreira AM, Johnson R, Jungreis I, Loveland J, et al. GENCODE reference annotation for the human and mouse genomes. Nucleic Acids Res. 2019;47(D1):D766–d73.

69. Martin M. Cutadapt removes adapter sequences from high-throughput sequencing reads. EMBnetjournal. 2011;17(1):10–2.

70. Aronesty E. Comparison of Sequencing Utility Programs. The Open Bioinformatics Journal. 2013;7(1):1–8.

71. Steinegger M, Söding J. MMseqs2 enables sensitive protein sequence searching for the analysis of massive data sets. Nat Biotechnol. 2017;35(11):1026–8.

72. Jumper J, Evans R, Pritzel A, Green T, Figurnov M, Ronneberger O, et al. Highly accurate protein structure prediction with AlphaFold. Nature 2021 596:7873. 2021;596(7873):583–9.

73. Varadi M, Bertoni D, Magana P, Paramval U, Pidruchna I, Radhakrishnan M, et al. AlphaFold Protein Structure Database in 2024: providing structure coverage for over 214 million protein sequences. Nucleic Acids Res. 2024;52(D1):D368–d75.

74. Akdel M, Pires DEV, Pardo EP, Jänes J, Zalevsky AO, Mészáros B, et al. A structural biology community assessment of AlphaFold2 applications. Nat Struct Mol Biol. 2022;29(11):1056–67.

75. Rost B, Sander C. Conservation and prediction of solvent accessibility in protein families. Proteins. 1994;20(3):216–26.

