## Supplementary Material for "A complete map of human cytosolic degrons and their relevance for disease"


**Fig. S1.** *Comparison of the five sublibraries by flow cytometry*. p.2
**Fig. S2.** *Effect of inhibiting autophagy and cullin-RING ligases.* p.3
**Fig. S3** *Transformation of degron scores to abundance scores & definition of degron strength.* p.4
**Fig. S4.** *Effect of homodipeptide positioning within tiles.* p.5
**Fig. S5.** *Comparison of polyalanine positioning within tiles.* p.6
**Fig. S6.** *The average degron score of tiles ending in each amino acid residue.* p.7
**Fig. S7.** *Lysine residues near the C-terminus counter degrons and increase protein abundance.* p.8
**Fig. S8.** *Previously characterized degron motifs span a wide range of degron scores.* p.9
**Fig. S9.** *Correlations with average tile exposure*. p.10
**Fig. S10.** *KRTAP11-1 contains highly exposed and disordered degrons.* p.11
**Fig. S11.** *Lasso regression analysis.* p.12
**Fig. S12.** *Correlation of PAP scores with validation tile GFP:mCherry measurements.* p.13
**Fig. S13.** *Correlations between ΔPAP and abundance scores in exposed and buried regions.* p.14
**Fig. S14.** *Degron disrupting missense mutations increase the abundance of full-length LAP3.* p.15
**Fig. S15.** *Abundance of full length ASPA variants correlates with the tile abundance.* p.16
**Fig. S16.** *ΔPAP predicts the abundance of ASPA variants in the 256-260 loop.* p.17
**Fig. S17.** *ΔPAP predicts the abundance of missense variants in exposed regions*. p.18
**Fig. S18.** *FACS gating strategy.* p.20
**Table S1.** *Mutations to avoid NotI and BsiWI restriction sites.* p.21
**References** p.22

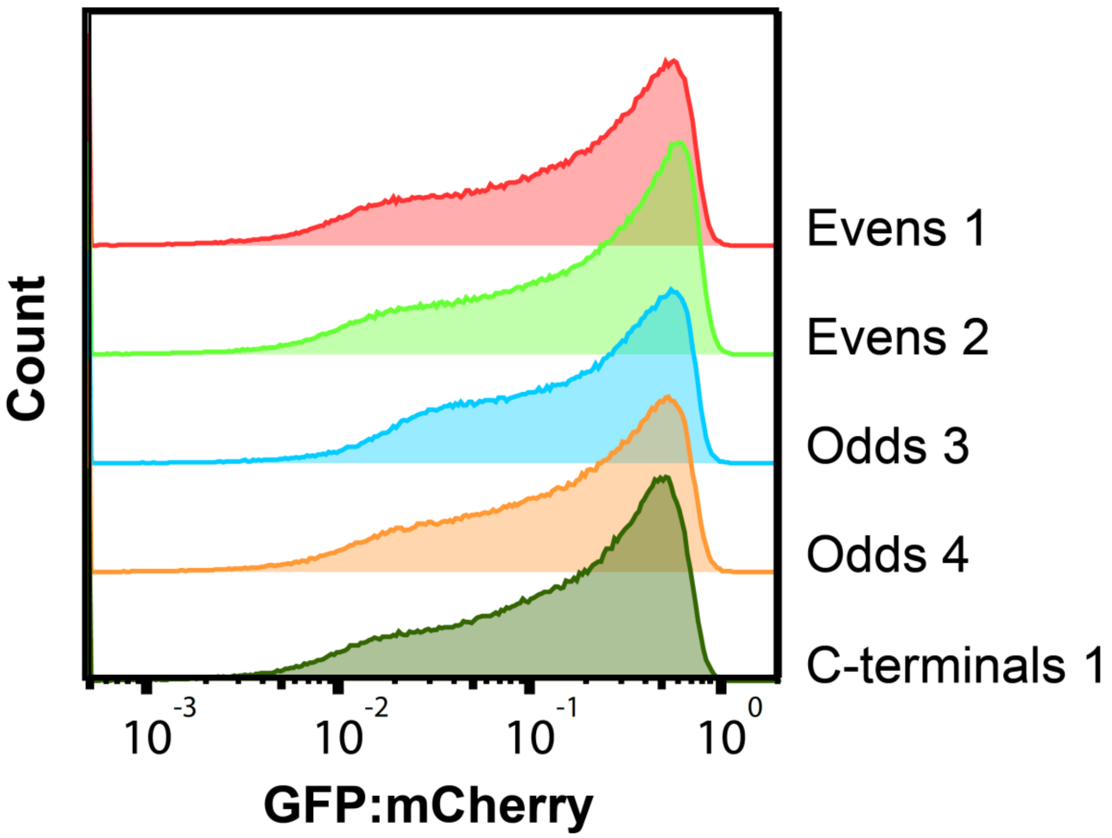


**Fig. S1 – Comparison of the five sublibraries by flow cytometry.**
Representative flow cytometry GFP:mCherry profiles of the five sublibraries (named evens 1, evens 2, odds 3, odds 4 and C-terminals 1) that were assayed. (Evens 1: n = 432,000, Evens 2: n = 451,000, Odds 3: 395,000, Odds 4: 415,000, C-terminals: n = 467,000).

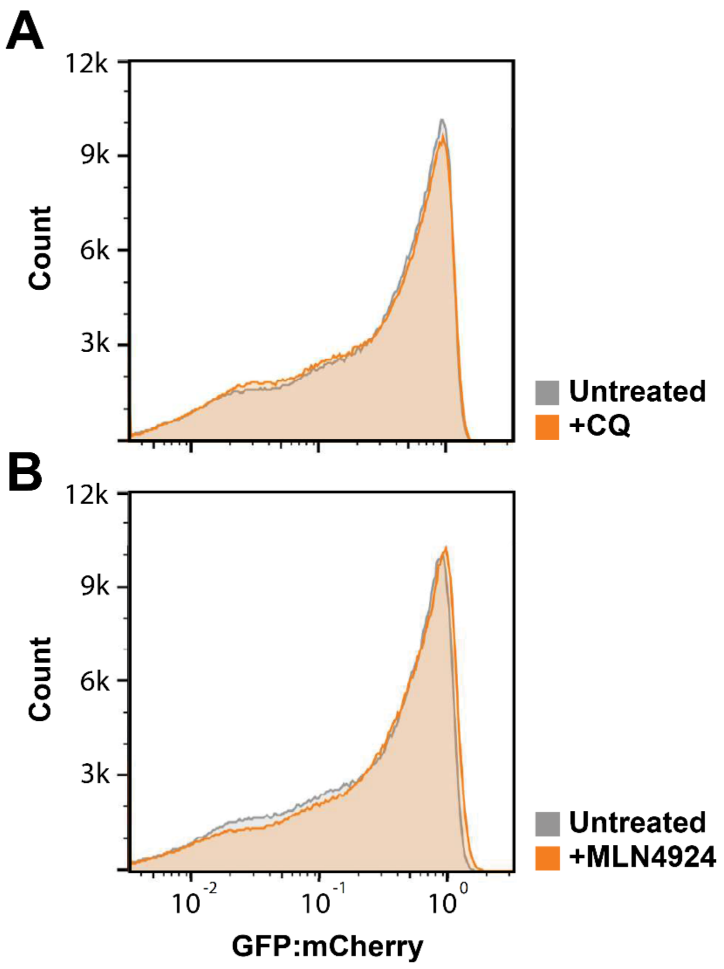


**Fig. S2 – Effect of inhibiting autophagy and cullin-RING ligases.**
Representative flow cytometry GFP:mCherry profiles of the Evens 1 sublibrary treated (orange) or not treated (grey) with (A) 20 µM chloroquine (CQ) (n = 768,530) or (B) the NEDD8 E1 activating enzyme inhibitor MLN4924 (n = 800,000).

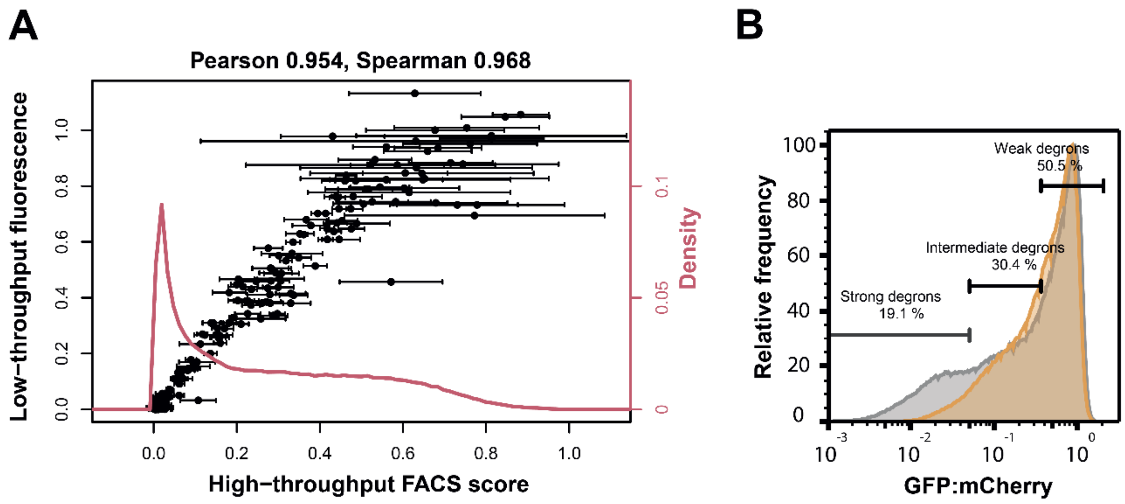


**Fig. S3 – Transformation of degron scores to abundance scores and definition of degron strength**. (A) The abundance scores (x-axis) are made by transforming degron scores to the GFP/mCherry fluorescence distribution measured by flow cytometry (red line). The validation peptides (black dots) were measured using flow cytometry in low throughput and correlate linearly with the high abundance scores on this scale. The error bars are the degron score standard deviations propagated using the derivative of the transforming function. (B) The degrons within the tiles were categorized into strong, intermediate and weak degrons based on their position within the FACS distribution. The ones within the peak that was not affected by BZ treatment (15 μM for 16 h) were defined as weak degrons, the ones with a GFP:mCherry ratio lower than the lowest 5% of the BZ were defined as strong degrons and the rest as intermediate.

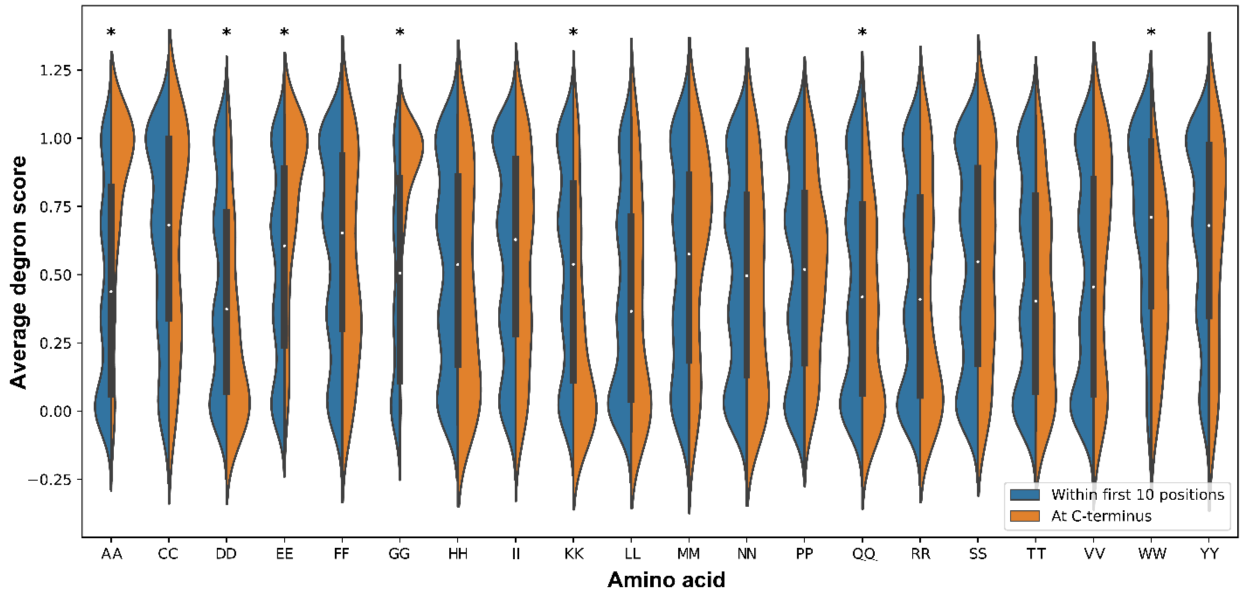


**Fig. S4 – Effect of homodipeptide positioning within tiles.**

Violin plots showing the degron score distribution of tiles that contain a dipeptide consisting of two of the same amino acid residues for each amino acid, when the homodipeptide is within the ten N-terminal positions (blue) or at the last two positions in the C-terminus (orange). The white dot is at the median of the two populations and the bars indicate the interquartile range (IQR). An asterisk indicates statistically significant differences between the two populations based on Mann-Whitney U test after Bonferroni correction ($p < 0.05/20$).

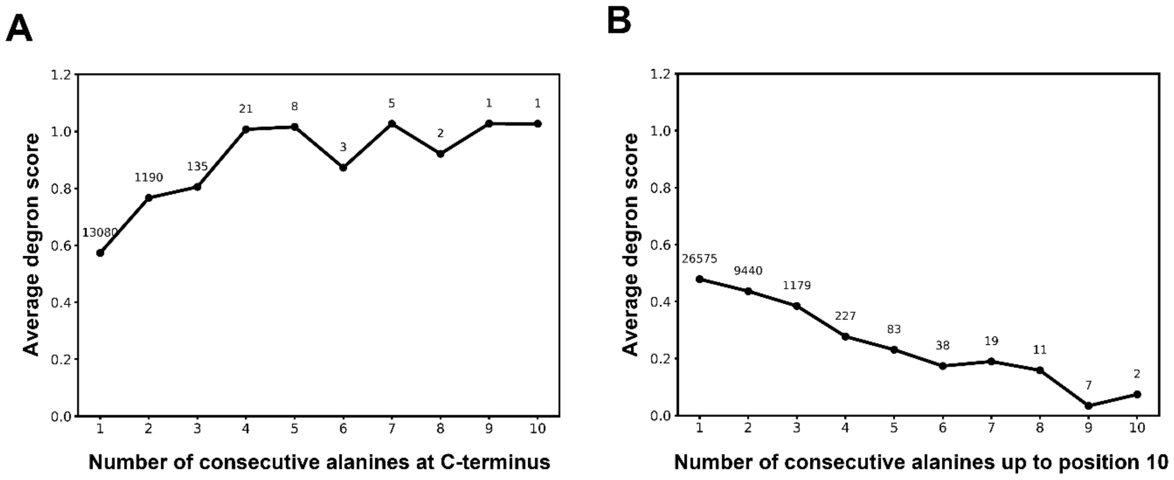


**Fig. S5 – Comparison of polyalanine positioning within tiles.**

Line plot showing the average degron score (y-axis) of tiles that have the indicated number of consecutive alanine residues (x-axis) when (A) at the C-terminus and (B) within the first ten positions of the tiles. The number of tiles used to calculate each average degron score is given above the data points.

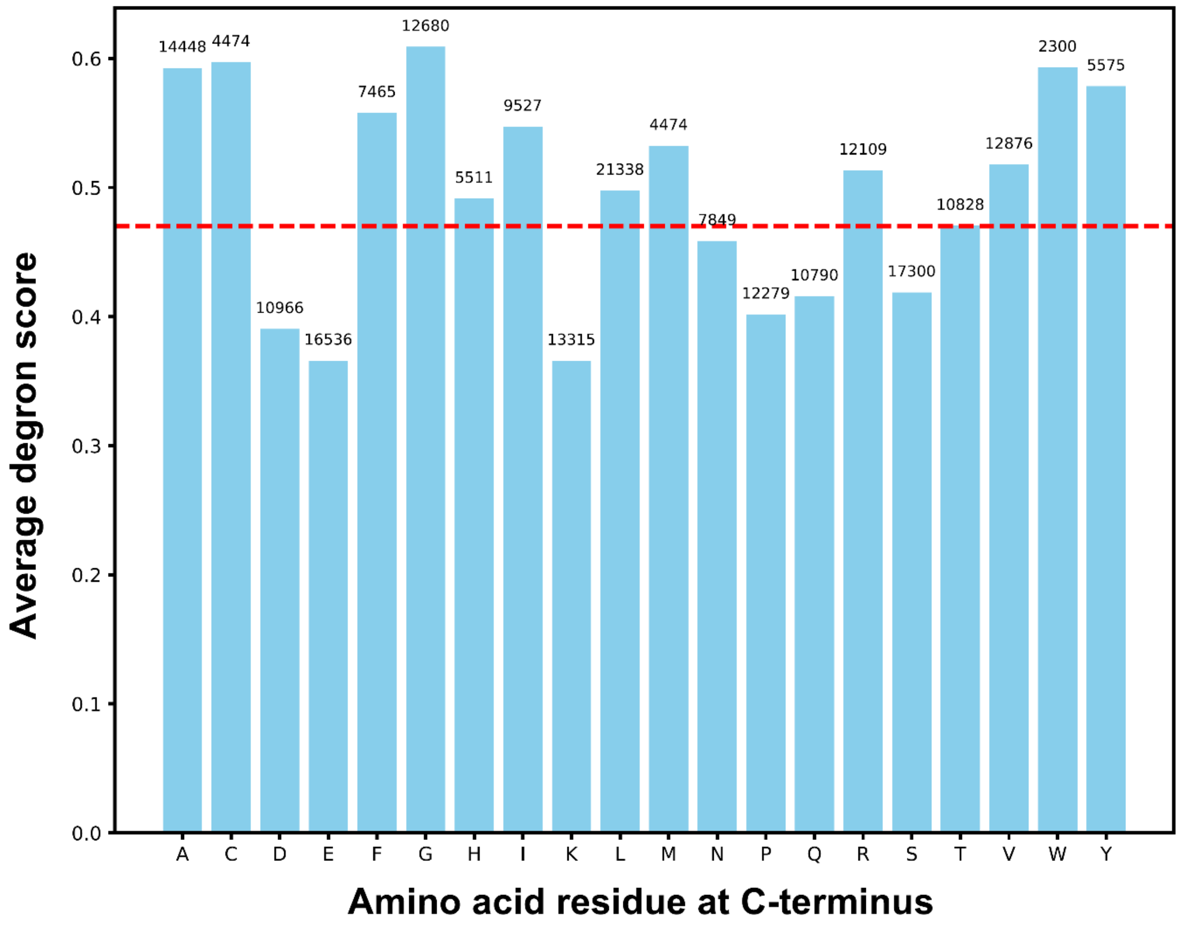


**Fig. S6 – The average degron score of tiles ending in each amino acid residue.**
Plot showing the average degron potency of tiles ending in each of the amino acids. Note that histidine, asparagine and threonine are the amino acids that have the closest to a neutral effect, given that the average degron score of all the tiles is 0.48 (red line). The number of tiles used to calculate each average degron score is given above the bars.

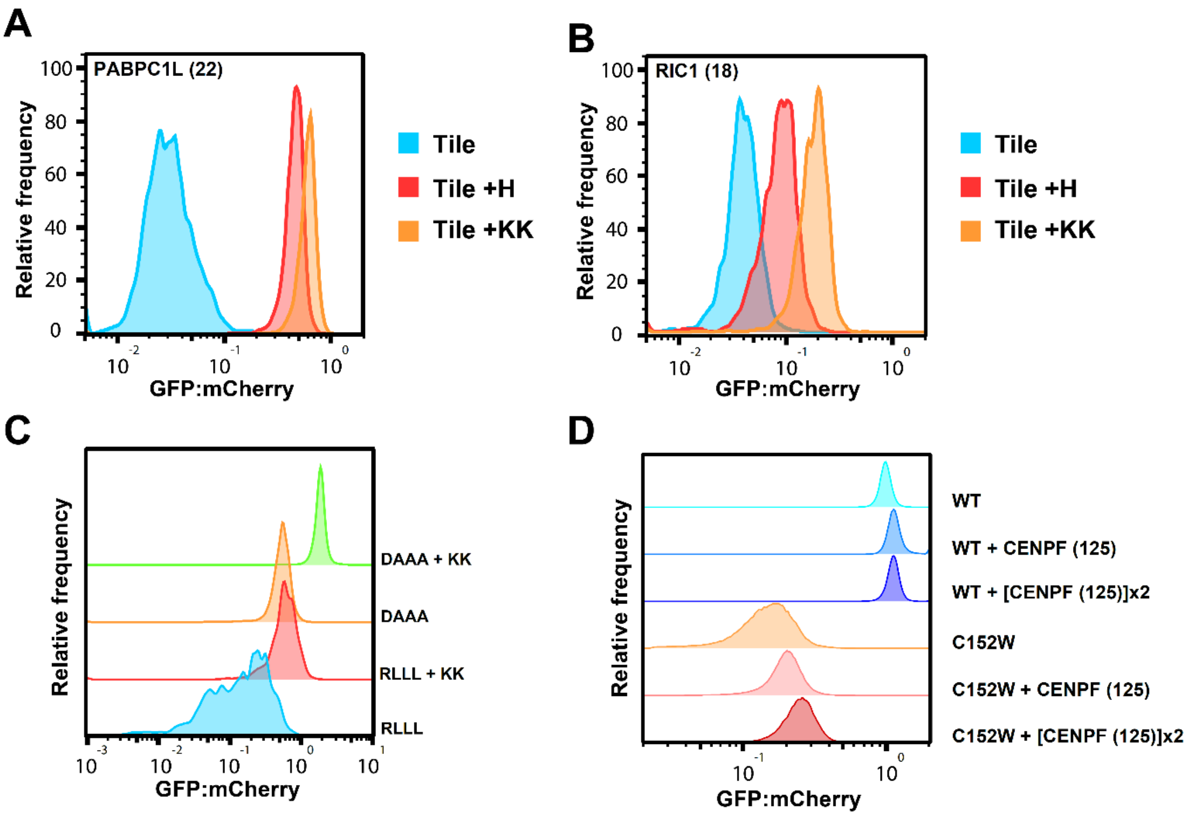


**Fig. S7 – Lysine residues near the C-terminus counter degrons and increase protein abundance.** (A) Representative flow cytometry GFP:mCherry profiles of PABPC1L tile 22 (PYGVITSAKVMTEGGHSKGFGFVCFSSPEE) and (B) RIC1 tile 18 (VAVNNKYRLMAFGCVSGSVQVYTIDNSTGA) with the C-terminal degrons -EE* and -GA* respectively. The profiles of each tile sequence (PABPC1L: n = 1312, RIC1: n = 1931) and of the sequence C-terminally fused with histidine (PABPC1L: n = 1749, RIC1: n = 1720) or a lysine dipeptide (PABPC1L: n = 1622, RIC1: n = 1684) are shown. (C) Representative flow cytometry GFP:mCherry profiles of the APPY peptide (CALLQS**RLLL**SAPRRAAATARY) and the DAAA mutant (CALLQS**DAAA**SAPRRAAATARY) with (RLLL: n = 1670, DAAA: n = 1557) or without (RLLL: n = 1661, DAAA: n = 1603) a C-terminal lysine dipeptide extension. (D) Representative flow cytometry GFP:mCherry profiles of the unstable ASPA C152W variant and ASPA WT with no fused C-terminal sequence (C152W: n = 42,841, WT: n = 135,835), with one C-terminally fused CENPF tile 125 (ETSEGLNSDLEMHADKSSREDIGDNVAKVN) (C152W: n = 119,579, WT: n = 69,444) or with two of the same tile in tandem (C152W: n = 48,512, WT: n = 182,700). All experiments were performed in duplicate.

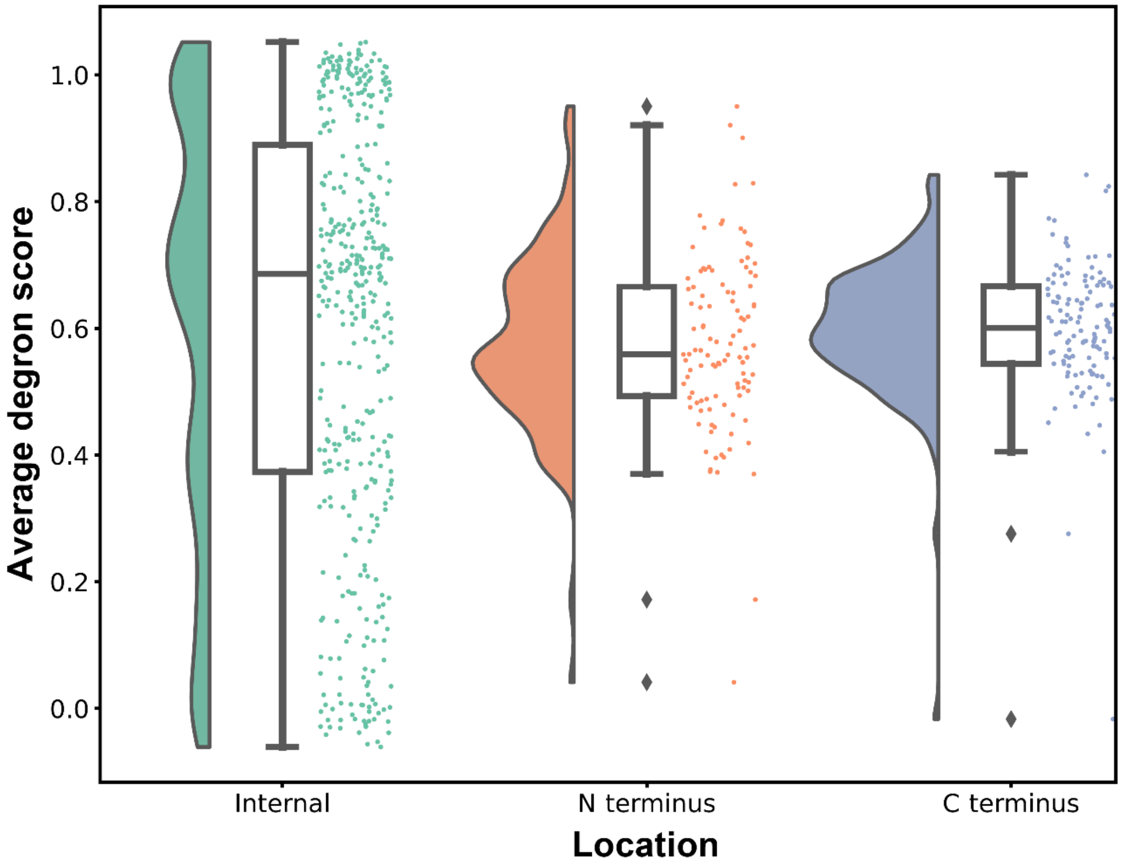


**Fig. S8 – Previously characterized degron motifs span a wide range of degron scores.**
Average degron score of tiles containing previously described degron motifs that are internal, at the N- or at the C-terminus (1). Specific motifs and their average scores are included in the source data file.

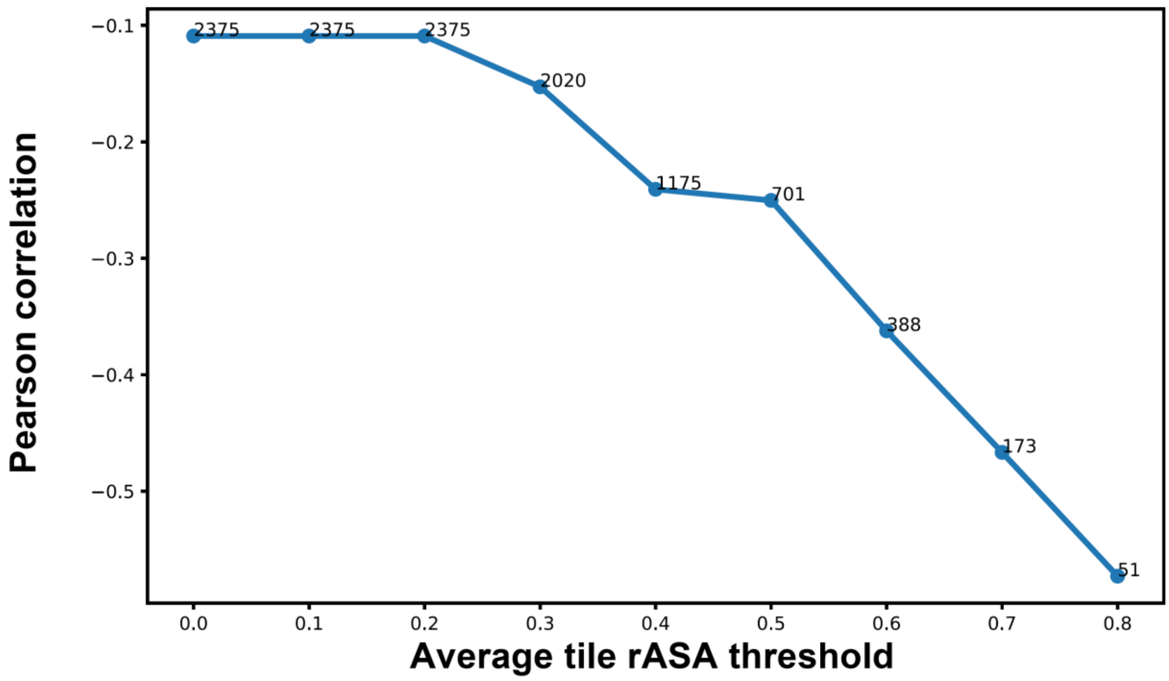


**Fig. S9 – Correlations with average tile exposure.**
Line plot of the Pearson correlations between protein stability index and degron score when considering proteins with an increasing average tile rASA. The number of proteins included in the calculations is indicated.

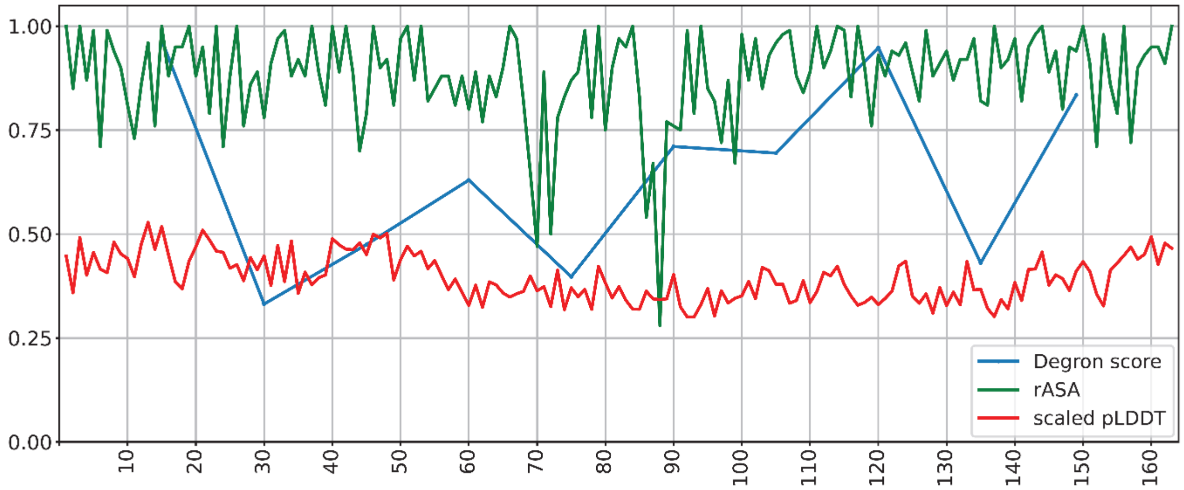


**Fig. S10 – KRTAP11-1 contains highly exposed and disordered degrons.**
Line plot showing the degron map (scores of degrons in the proteins) of KRTAP11-1 (blue line). The rASA (green line) and pLDDT (red line) of each residue of the protein are also shown.

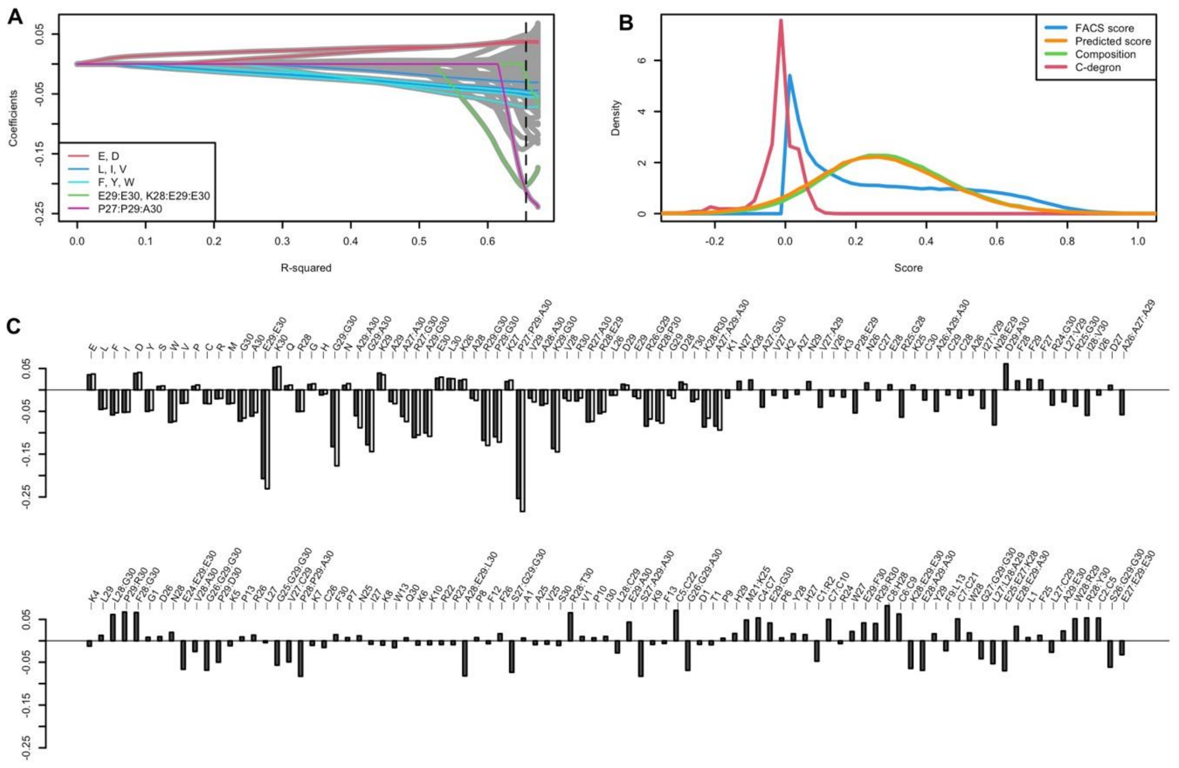


**Fig. S11 – Lasso regression analysis**.

(A) Lasso pathway showing the magnitude of individual feature coefficients (lines) as a function of the fraction of R-squared. Lasso regularization is strongest to the left and as the regularization is relaxed, coefficients increase and more become non-zero to make a better model (higher R-squared) until the point where increasing the coefficients no longer improves the model. The dashed line shows the regularization strength that yields the maximum number of 179 significant features in a non-regularized linear model (Pearson 0.81, RMSE 0.137, dark bars in panel C). (B) Distribution of predicted scores from the "Human30" model trained on selected features without regularization (Pearson 0.80, RMSE 0.140, light bars in panel C) (see Methods). (C) Bar plot showing the coefficients of the two non-regularized linear models (dark and light bars) ordered according to feature importance (see Methods). Several pair features and most of the triplets seem to account for non-linearities rather than representing individual motifs. For example, A27, A29, A30 are all destabilizing individually and if combined in pairs (A27:A30, A29:A30) or triplets (A27:A29:A30), an additional destabilization is added because the combinations are stronger than the sums. Similarly, the strength and early appearance of the E29:E30 C-degron indicates that this is important but also abundant, in fact the most abundant pair of the considered, and thus this is able to explain more of the data. However, it decreases in magnitude around R-squared 0.66 likely due to the activation of the triplet coefficient K28:E29:E30 that explain a subset of the same peptides and is identified as a stronger effect (panel A, green lines).

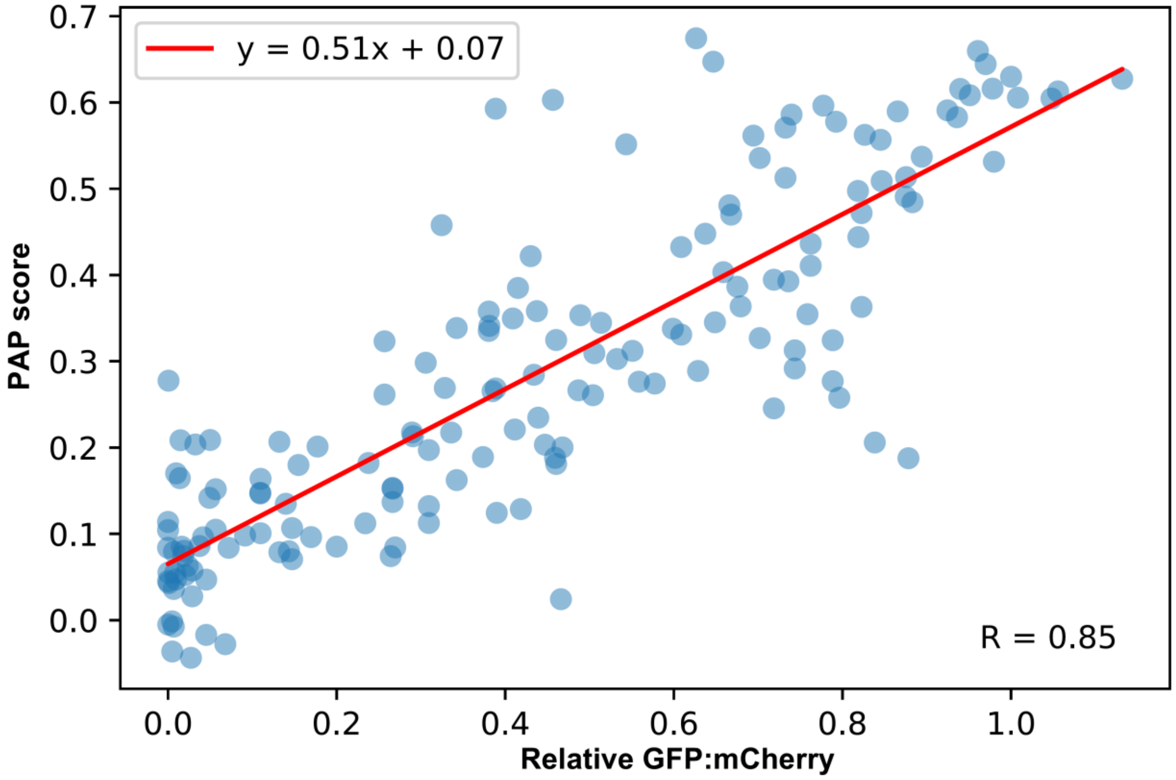


**Fig. S12 – Correlation of PAP scores with validation tile GFP:mCherry measurements**. Correlation of the low-throughput measurements of GFP:mCherry with their equivalent PAP scores shows the ability of PAP to predict experimental measurements instead of assay scores.

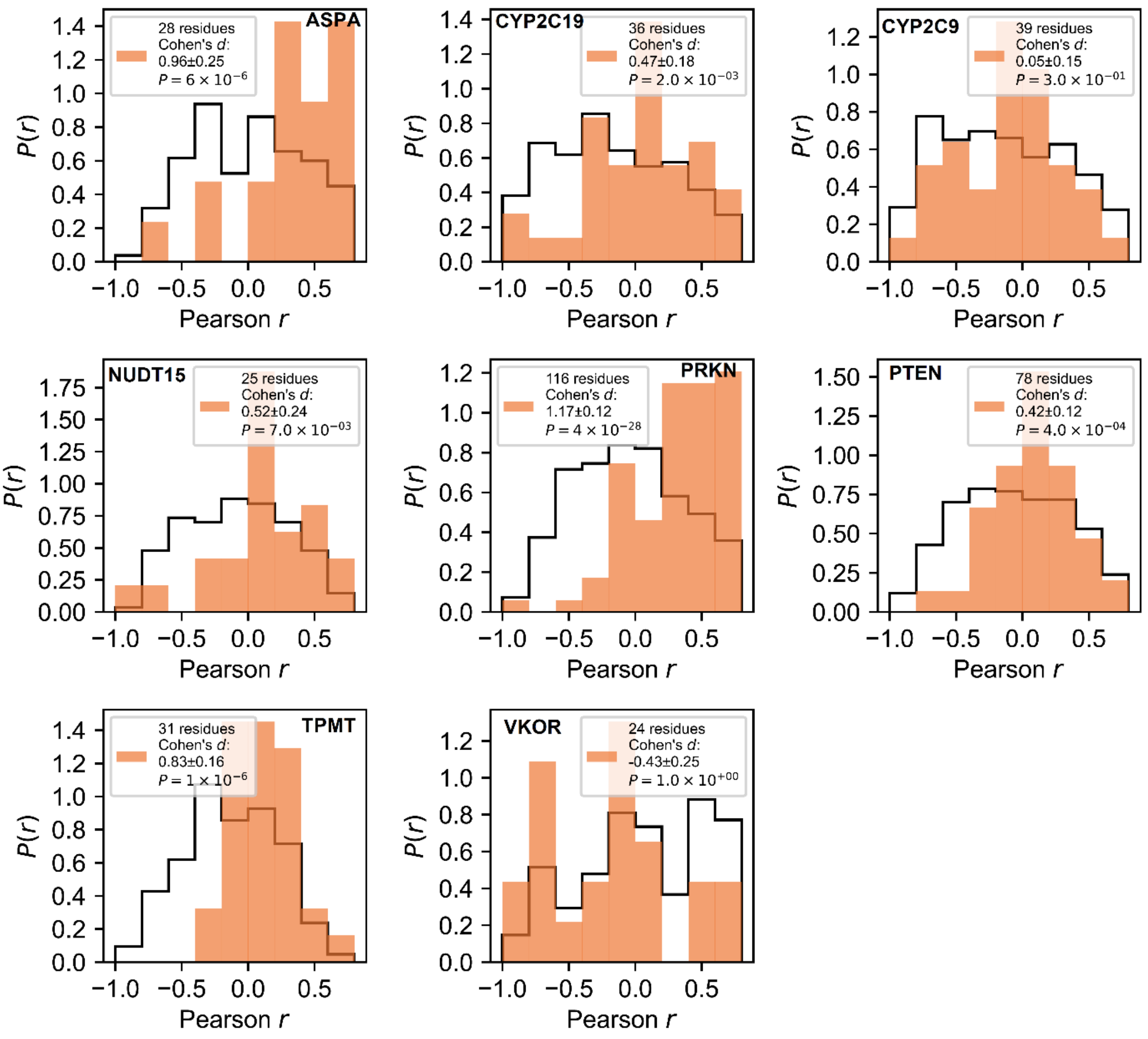


**Fig. S13 – Correlations between ΔPAP and abundance scores in exposed and buried regions.** Histograms of Pearson correlation coefficients for two subsets of data (rASA > 0.7 and rASA ≤ 0.7). The histograms (black lines) show the density distribution of Pearson correlation coefficients for residues with rASA ≤ 0.7, while the bar plots (orange bars) display the equivalent density distribution for residues with rASA > 0.7. Cohen's d, representing the effect size, and the p-value are annotated on each plot. Bootstrapping with 10,000 resamples was used to estimate the confidence interval of Cohen's d and to compute a p-value based on the Brunner-Munzel test statistic. Note that the Pearson correlation r between ΔPAP and abundance scores is predominantly negative for exposed regions (rASA > 0.7), while it tends to be close to zero or positive for less exposed regions (rASA ≤ 0.7). In all cases except CYP2C9 and VKOR, which both have extended regions that are buried in the membrane, there is a significant difference in the Pearson correlation between exposed and less exposed residues.

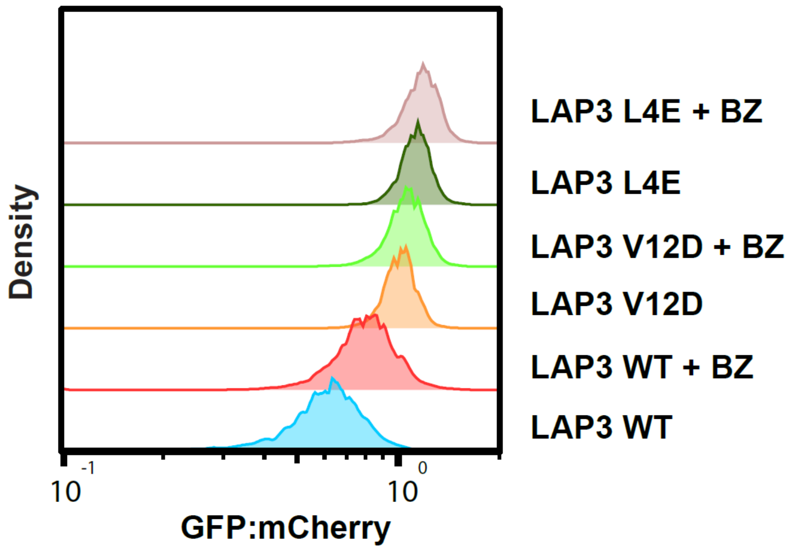


**Fig. S14 – Degron disrupting missense mutations increase the abundance of full-length LAP3.** Representative flow cytometry GFP:mCherry profiles of the LAP3 WT and of two missense mutations (L4E and V12D) predicted to increase peptide abundance (ΔPAP L4E = 0.09 ΔPAP V12D = 0.08, tile size = 15). The equivalent profiles of BZ (15 µM for 16h) treated cells are also shown (WT: n = 6,323, WT + BZ: n = 5,433, V12D: n = 5,489, V12D + BZ: n = 5,350, L4E: n = 5,450, L4E + BZ: n = 5,174). The experiment was performed in duplicate.

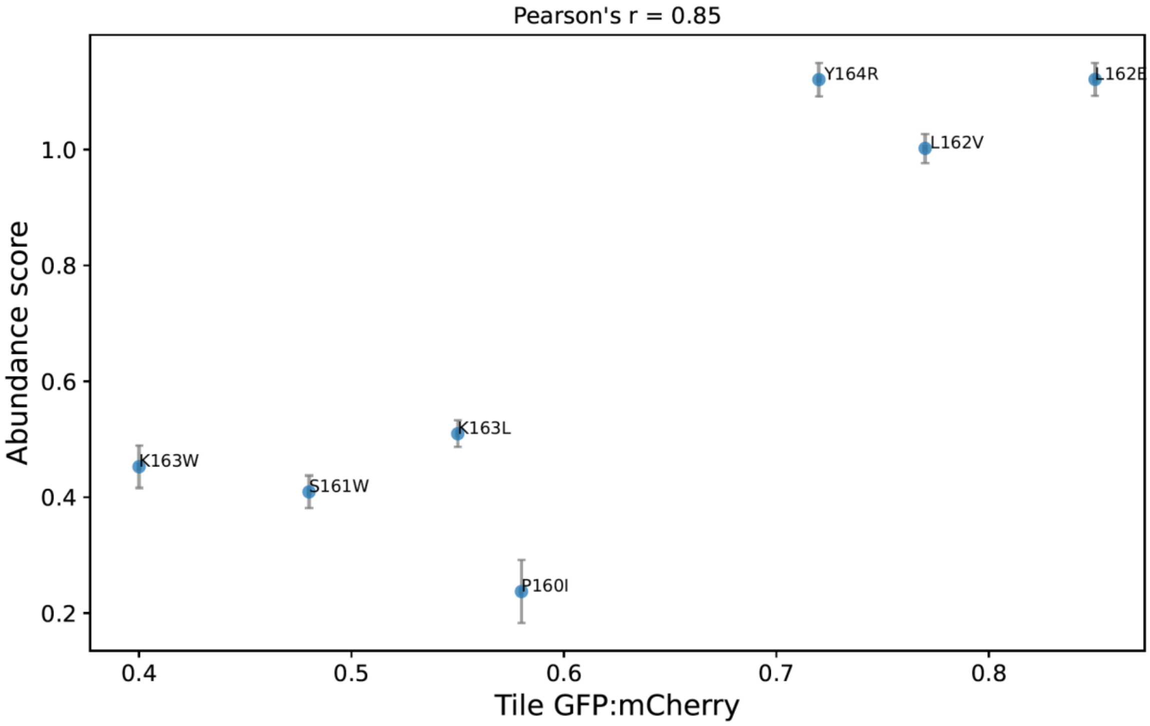


**Fig. S15 – Abundance of full length ASPA variants correlates with the tile abundance.** Correlation plot of eight-residue long ASPA tile covering the positions 159-166 and containing the annotated mutations with the equivalent full-length ASPA variants. The vertical error bars show the standard deviation of the abundance score of each ASPA variant and the horizontal show the standard error of the GFP:mCherry measurements. The experiment was performed in duplicate.

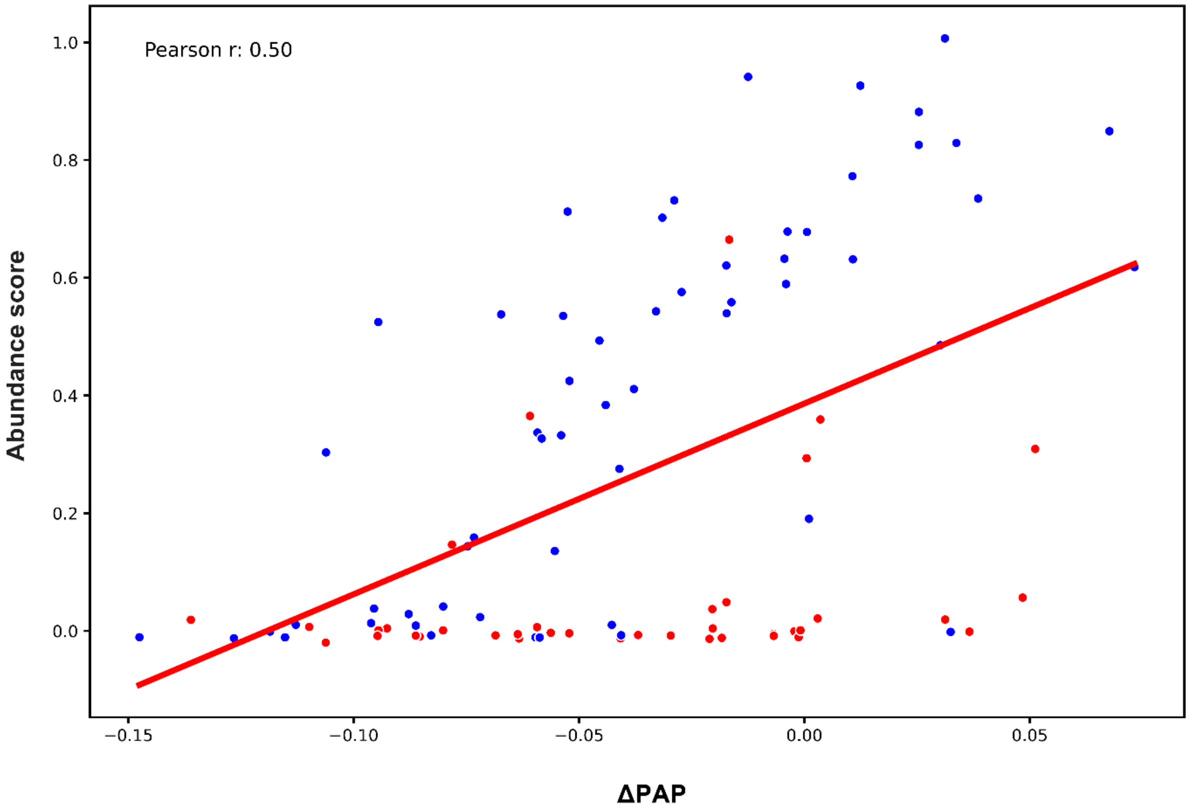


**Fig. S16 –ΔPAP predicts the abundance of ASPA variants in the 256-260 loop.**
Correlation plot of ΔPAP with abundance scores in the ASPA loop consisting of residues 256-260. Note that the correlation is strong for most substitutions in all the positions except for residues P257 and P260 (red points), potentially indicating structurally important prolines for the loop.

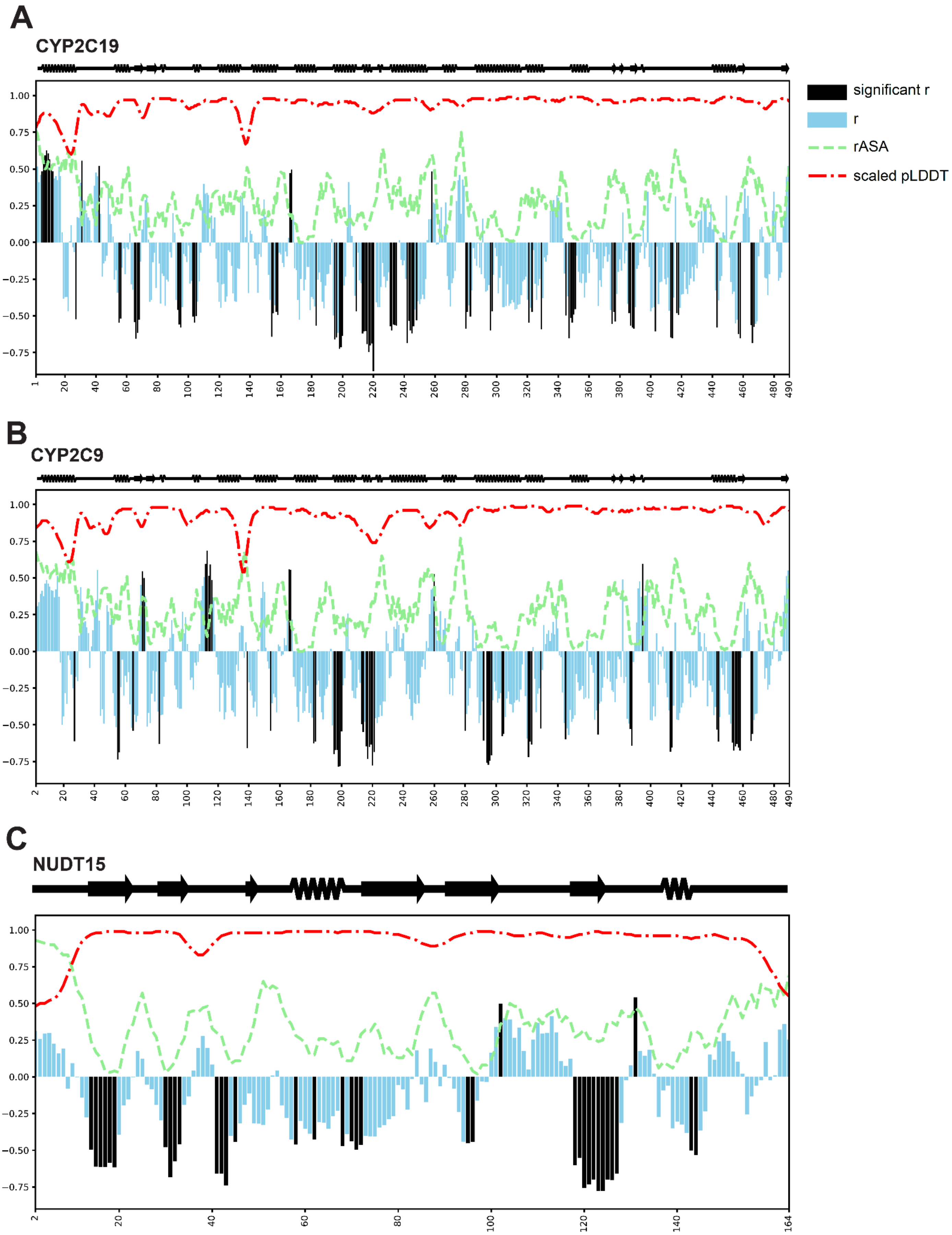
A
CYP2C19
significant r
r
rASA
scaled pLDDT
B
CYP2C9
C
NUDT15

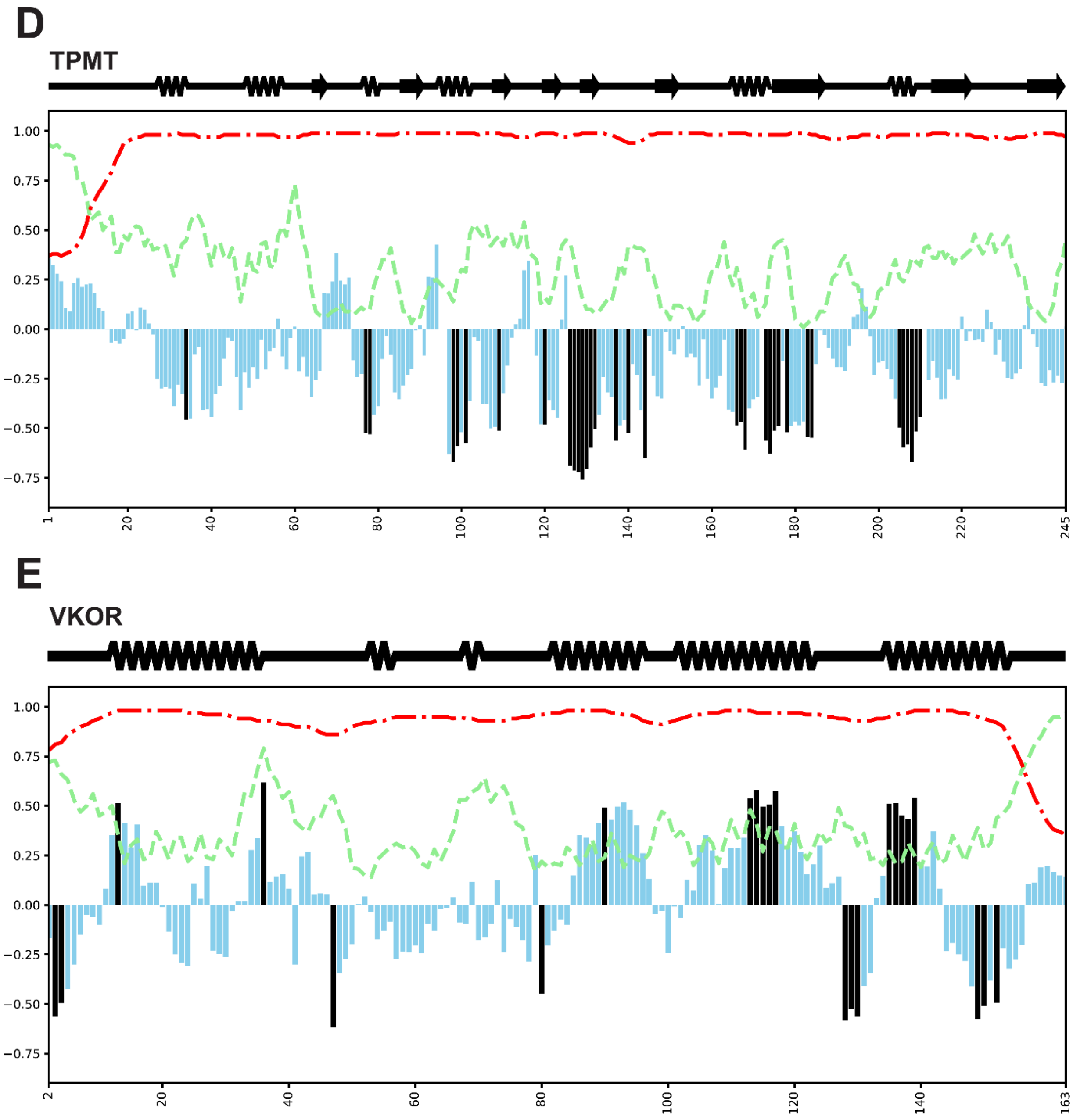


**Fig. S17 – ΔPAP predicts the abundance of missense variants in exposed regions.**
Correlation map ΔPAP against abundance score of single amino acid substitution variants of (A) CYP2C19, (B) CYP2C9, (C) NUDT15, (D) TPMT and (E) VKOR. Bars show the Pearson's correlation of all the scored variants against their predicted ΔPAP for a sliding window of five residues. A t-test for Pearson's correlation coefficient was performed for every five-residue window. Black bars indicate statistically significant correlations after Bonferroni correction with $p < 0.05/m$, where m is the number of tests conducted for each protein. The average rASA (green line) and scaled pLDDT (red line) of each residue window are also shown. The secondary structure of each protein is shown above each plot. Note that a good correlation is observed mostly in unstructured exposed regions. This applies even to some VKOR loops that face the lumen of the endoplasmic reticulum.

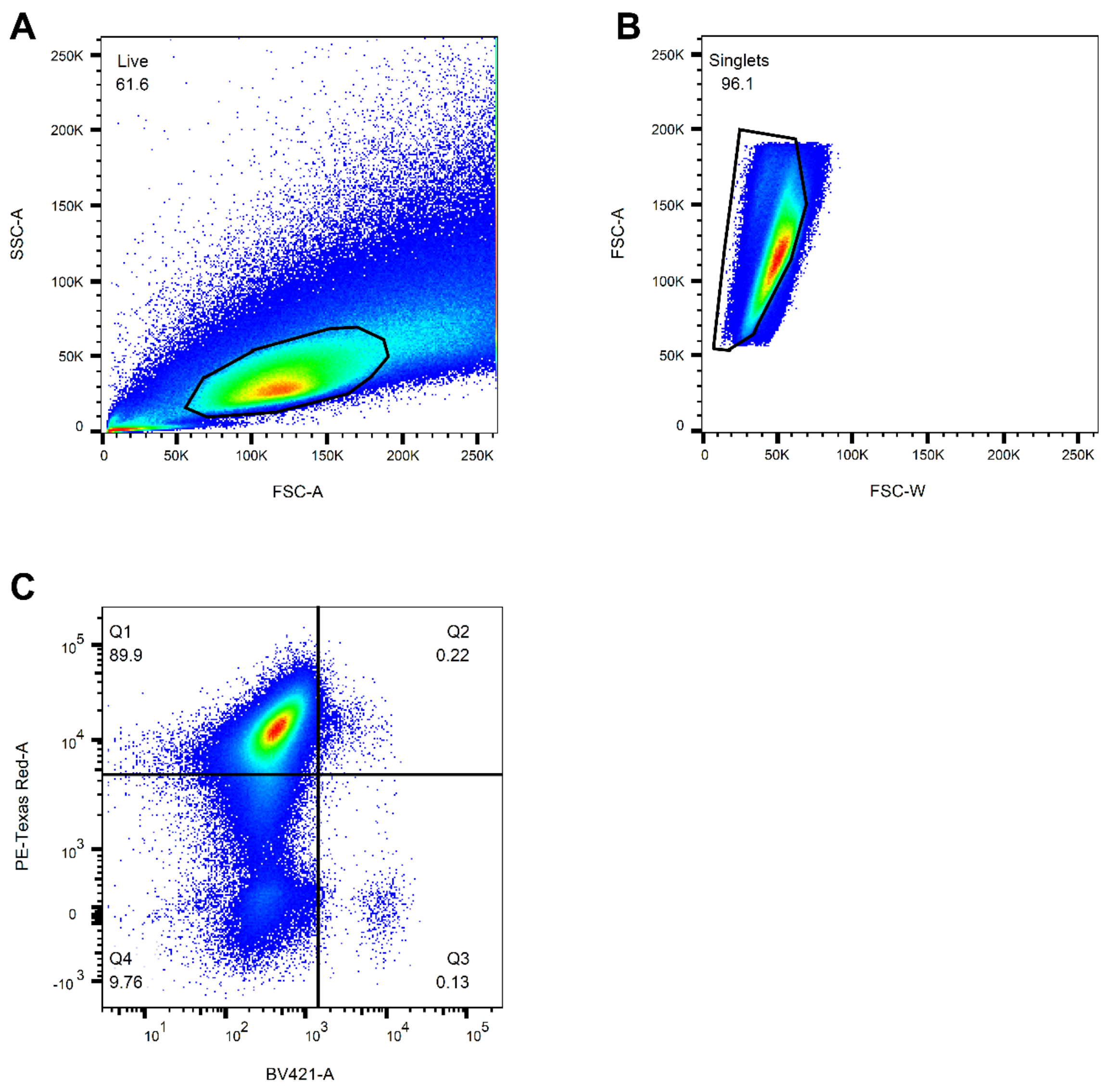


**Fig. S18 –FACS gating strategy.**
(A) Gating of live population based on side (SSC) and forward scatter (FSC) (n = 886,000). (B) Gating of single cells (singlets) based on the forward scatter pulse area (FSC-A) and width (FSC-W) (n = 546,000). (C) Gating of the BFP (BV421-A) negative and mCherry (PE-TexasRed) positive recombinant population of cells (n = 524,000).

**Table S1**

*Mutations to avoid NotI and BsiWI restriction sites*

| Enzyme | substrate | occurrences | frame1subst | frame2subst | frame3subst |
|---|---|---|---|---|---|
| NotI | GCGGCCGC | 677 | GCaGCCGC | GCGcCCGC | GCGGaCGC |
| BsiWI | CGTACG | 389 | CGaACG | CGTtCG | CGTAtG |

### References

1. Szulc NA, Stefaniak F, Piechota M, Soszyńska A, Piórkowska G, Cappannini A, et al. DEGRONOPEDIA: a web server for proteome-wide inspection of degrons. Nucleic Acids Res. 2024.
